# Microbial Life Inside *Posidonia* Seeds: Beneficial Endophytes and Implications for Marine Plant Health

**DOI:** 10.1101/2025.11.19.689238

**Authors:** Dalila Crucitti, Alberto Sutera, Roberto De Michele, Francesco Carimi, Stefano Barone, Fabio Badalamenti, Davide Pacifico

**Affiliations:** Institute of Biosciences and Bioresources (IBBR), CNR., Via Ugo La Malfa 153, 90146 Palermo, PA, Italy; Department of Agricultural, Food and Forest Sciences (SAAF), University of study of Palermo, Viale delle Scienze, Palermo 90128, Italy; Institute for the study of Anthropic Impacts and Sustainability of the Marine Environment (IAS), CNR, Lungomare Cristoforo Colombo 4521,90149, Palermo, Italy

**Author notes:** (F.C.); (D.P.). (S.B.). (F.B.).

**Keywords:** seagrass, *Posidonia oceanica*, endophytes, marine microbiota, *Plant growth promoting bacteria*, *Halophytophthora*, *Celerinatantimonas*, *Marinomonas*, *Vibrio*, *Kocuria*

## Abstract

Plant–microbe interactions are key drivers of plant health and ecosystem functioning, yet their roles in marine environments remain poorly understood. The seagrass *Posidonia oceanica*, a foundation species in the Mediterranean Sea, forms complex associations with microbial communities that influence its development and stress tolerance. Here, we provide the first evidence of culturable bacterial and fungal endophytes inhabiting *P. oceanica* seeds collected from central Mediterranean, a region representing a major center of the species’ genetic diversity. Using two different marine culture media, we isolated a diverse assemblage of endophytes, predominantly affiliated with *Marinomonas, Celerinatantimonas, Vibrio, Halomonas, Kocuria, Bacillus, Metabacillus, Lysobacter*, and *Aureimonas*, along with the fungi *Paecilomyces maximus* and *Halophytophthora* sp. Most bacterial isolates displayed plant growth–promoting (PGP) traits such as indole-3-acetic acid production and nitrogen fixation, supporting their potential contribution to seed germination and early seedling establishment. The detection of *Candidatus* Celerinatantimonas neptuna, a nitrogen-fixing symbiont previously described in *P. oceanica* roots, suggests a possible route of vertical transmission. Although fungal endophytes were less frequent, their presence indicates that *P. oceanica* seeds may serve as a reservoir of both beneficial and potentially pathogenic taxa. These findings expand our understanding of the *P. oceanica* holobiont, highlight the role of seeds in the persistence and dissemination of endophytic communities and lay the groundwork for the biotechnological use of seed-associated microbes in marine plant restoration and conservation, and in crop stress tolerance.

## 1. Introduction

Several species of endophytic bacteria, actinomycetes, fungi, archaea, and protists colonize internal plant tissues, establishing mutualistic interactions that confer various benefits to their hosts (Compant et al. 2013; Hardoim et al. 2015). These positive effects on plant growth and resistance to biotic and abiotic stresses arise both directly (from microbial metabolic activities such as nitrogen fixation, phosphate solubilization, hormone production, and synthesis of biocides or antimicrobial compounds) and indirectly, through the modulation of plant physiology, nutrient balance, or by occupying the ecological niches of pathogens, thereby enhancing stress tolerance (Compant et al. 2013; Santoyo et al. 2016; Khare et al. 2018; Pacifico et al. 2019; Ali et al. 2022). The most advanced studies on plant–microorganism interactions have revealed a wide range of potential biotechnological applications for crop improvement and environmental protection or restoration (Compant et al. 2019).

Plant endophytes can be transmitted horizontally through the environment (via soil or between plants), vertically through seeds and pollen, or via a mixed transmission mode (Frank et al. 2017; Nelson 2018). Vertical transmission ensures the inheritance of beneficial endophytes exhibiting plant growth-promoting (PGP) traits in seedlings, contributing to dormancy release, enhanced germination and growth, and protection against biotic and abiotic stressors (Truyens et al. 2015; Nelson 2018).

The Mediterranean seagrass *Posidonia oceanica* (L.) Delile is the dominant marine angiosperm in the Mediterranean Sea, where it forms extensive coastal meadows that support high biodiversity. These meadows provide crucial ecosystem services, including carbon sequestration, oxygen production, bioremediation, nursery habitat and shelter for fish, and protection against coastal erosion (Scanu et al. 2022). Over the past decade, however, the distribution and density of *P. oceanica* meadows have dramatically declined due to climate change and increasing human exploitation of coastal areas (Telesca et al. 2015; Scanu et al. 2022). Naturally, meadow expansion occurs slowly through the proliferation of lateral buds and vegetative fragment dispersal (Marbà 1998; Di Carlo et al. 2005). Long-distance dispersal depends on floating fruits carried by marine currents, which release large, fleshy seeds upon ripening (Ruocco et al. 2024). Under unfavorable conditions, large amounts of seeds are cast on beaches, where they rapidly lose viability, unless protected by fruit encasing or retrieved by tides (Sutera et al. 2024). Following disturbance events, *P. oceanica* grasslands exhibit a low natural recovery rate (Badalamenti et al. 2011; Castejón-Silvo and Terrados 2021) . Restoration programs have been initiated to recover degraded seagrass habitats through the transplantation of cuttings from donor meadows or seedling transplant (Calvo et al. 2021; Bacci et al. 2025). Although still limited by the availability of clonal material and human disturbances, these efforts have yielded encouraging results (Calvo et al. 2021; Bacci et al. 2025). A seed-based propagation strategy is more convenient than cutting transplant, and it preserves genetic diversity and existing meadows. In addition, a recent study developed a feasible protocol for long-term storage of *P. oceanica* seeds, allowing scalable propagule production and transplant (Sutera et al. 2025).

Endophytes are known to play key roles in nutrition and defense processes (Espinosa et al. 2010; Garcias-Bonet et al. 2016; Mohr et al. 2021; Torta et al. 2022). A recent study on the Australian seagrass *Halophila ovalis* revealed a complex endophytic microbiota associated with reproductive tissues (flowers, fruits, and seeds), including a core of bacterial species with PGP traits and nitrogen-fixing capabilities conserved across organs (Tarquinio et al. 2021). In *P. oceanica*, the endophytic microbial community has been characterized using molecular/metagenomic approaches (Garcias-Bonet et al. 2012, 2016) and culturing methods (Espinosa et al. 2010; Blanchet et al. 2017; Torta et al. 2022). Garcia-Bonet *et al*. (Garcias-Bonet et al. 2012) showed different compositions of *P. oceanica* endophytic community among plant organs (roots, rhizomes, and leaves). However, the presence of microbial endophytes in seeds remains unexplored.

In this study, we applied culturing techniques to isolate the endophytic microbiota associated with surface-sterilized *P. oceanica* seeds, with the aim of identifying vertically transmitted microorganisms potentially beneficial for seed germination and seedling development. Endophytic isolates from *P. oceanica* seeds could be exploited to extend the shelf life of propagation material used in future restoration programs. Furthermore, the availability of culturable microbial isolates represents an important first step toward elucidating the complex interactions between endophytes and marine angiosperms, and supports the development of novel model systems designed to simulate plant–microbe interactions under marine conditions at the laboratory scale. Culturable endophytes isolated from plants growing in extreme environments, such as seagrasses, may also present biotechnological potential to enhance stress tolerance to crops.

## 2. Materials and Methods

### 2.1. Plant Material Collection and sterilization

From half May to early June 2021, 44 *P. oceanica* seeds, still completely enclosed by fruits, brought naturally ashore by the current, were harvested in five distinct locations on the Sicilian coast: Cornino (38.093 N, 12.661 E) (10 seeds), Erice (38.046 N, 12.548 E) (5 seeds), and Marsala (37.740 N, 12.472 E) (9 seeds) along the western coast; San Nicola (38.009 N, 13.624 E) (8 seeds) in the northern coast, in the Tyrrhenian Sea; and Sciacca (37.504 N, 13.063 E) (12 seeds), in the southern coast, in the Strait of Sicily (Figure 1). The fruits were transported to the laboratory, washed in sterile artificial seawater (ASW, 3.8% salinity) to remove plant and sand residues, then stored individually for 24-48 h in 15 mL of ASW at 4°C before processing. Under aseptic conditions, fruits were dissected with a sterile blade to extract seeds. Each seed was surface disinfected to inactivate the micro-organisms present on the surface by sequential immersion for 2 min in 70% (v/v) ethanol, 3 min in 7% (v/v) sodium hypochlorite (10% active chlorine), 1 min in 70% (v/v) ethanol, followed by three rinses of 2 min in sterile distilled water (SDW).

**Figure 1.**
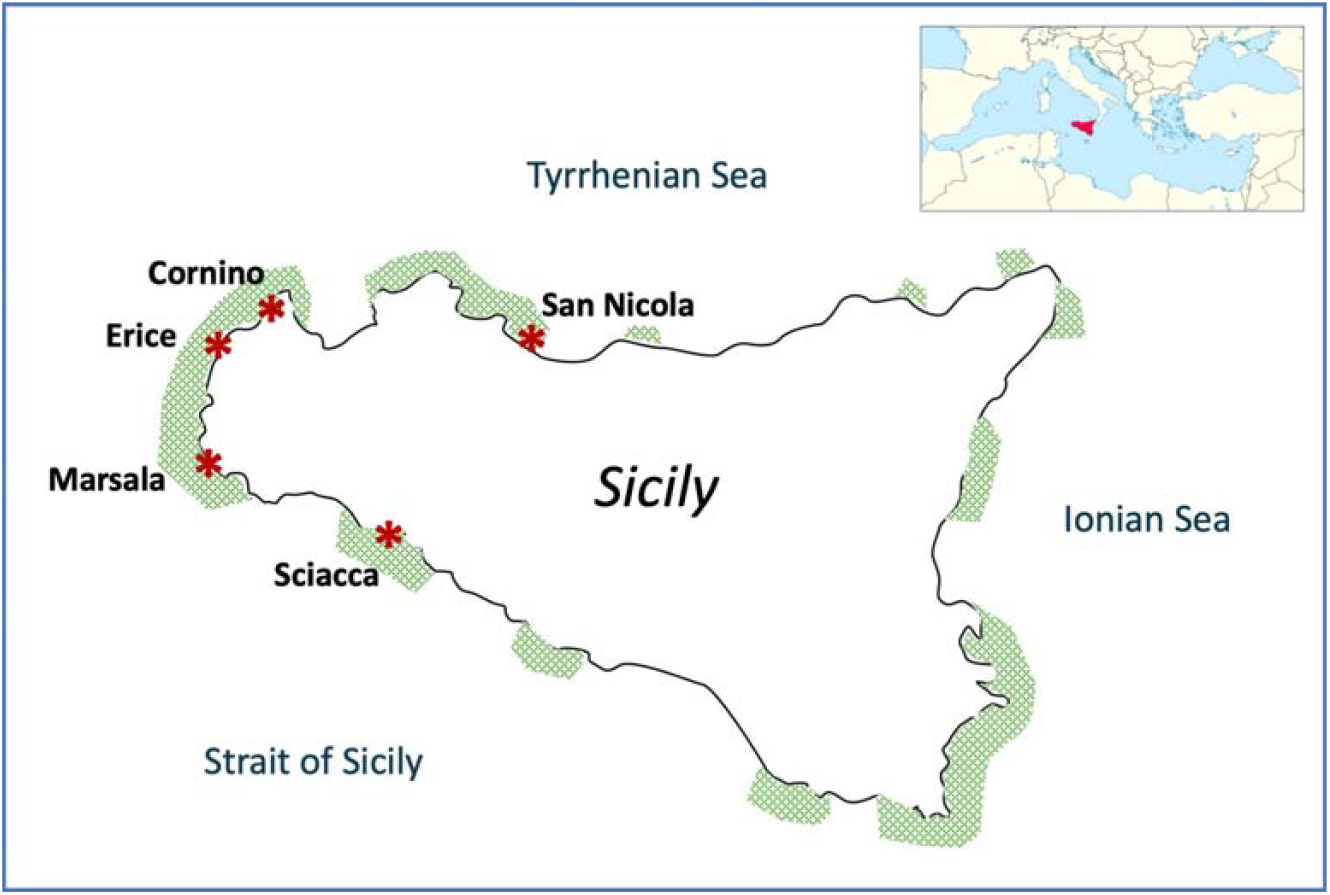
Locations, along the Sicilian coast, where cast fruits were collected. Shaded areas indicate the approximate current distribution of *P. oceanica* meadows (adapted from Calvo et al. 2010).

### 2.2. Endophyte isolation

Each surface-disinfected seed was weighed (Appendix 1) and longitudinally cut with a sterile blade. Half seed was then ground in 3 mL of SDW under aseptic conditions. Preliminary ten-fold serial dilution assays were performed on a subsample of ten seeds to assess the microbial load of the homogenates. Undiluted seed homogenate (100 µL) were plated on saline Nutrient Agar plates (NA: 3 g yeast extract, 5 g peptone, 20 g NaCl, 15 g Agar, 200 mL ASW, SDW up to 1 L, pH 6.8-7.2), and saline Starch Yeast extract Glucose Agar plates (SGY, 15 g starch, 5 g glucose, 5 g yeast extract, 20 g NaCl, 20 g Agar, 200 mL ASW, SDW up to 1 L, pH 6.8-7.2). All media were autoclaved for 20 min at 121 °C before use. To confirm surface disinfection, aliquots of the water used in the final rinse were plated onto the same media (negative control plates). Plates were then incubated at 26 °C in the dark and observed daily up to twenty days. Total colonies were counted, and the colony-forming unit (CFU)/g of fresh material was calculated for both media according to Silva *et al*. (da Silva et al. 2013) (Appendix 1).

Microbial colonies were initially grouped according to colony morphological features (shape, color, elevation, surface, margin, aerial mycelium, presence of exudate, and growth rate) and microscopic similarity (Appendix 2), and at least two representative colonies of each morphotype for each organ and host sample were selected and purified streaking agar plates. Pure bacterial colonies were cultured in 5 mL of the corresponding broth, incubated at 28 ± 2 °C for 24 h. Five hundred microliters were stored as pure cultures at −80 °C in glycerol (50% v/v). The remaining volume of the cell culture was pelleted by centrifuge at 2,600 rcf for 10 minutes and used for DNA extraction.

### 2.3. Endophyte molecular identification

Molecular identification was performed through sequence analysis of PCR-amplified bacterial 16S rDNA genes and fungal internal transcribed spacer (ITS1, 5.8S, ITS2) rDNA regions.

Bacterial pellets were resuspended in 500 µL of Tris-EDTA buffer (TE, pH 8.0) and enzymatically lysed with Proteinase K (20 mg/mL) for 60 minutes at 37 °C. Genomic DNA was purified using the phenol/chloroform/isoamyl alcohol (25:24:1) method (Wilson 1997) and resuspended in 100 µL TE. Bacterial endophytes were identified by sequencing ∼1500 bp of the 16S rDNA with the universal primers 27f and 1492r following the original PCR protocol (Lane 1991) using the DreamTaq DNA Polymerase (5 U/µL, Thermo Fisher Scientific).

Pure fungal cultures were obtained on SGY plates incubated for seven days at 28 ± 2 °C. The aerial mycelium (200 µg) of each selected fungal colony was scraped, placed in clean tubes and disrupted with tungsten carbide beads in a Tissue Lyser II (Qiagen) set at 30 Hz for 2 min. Fungal DNA was extracted following a Cetyl trimethyl ammonium bromide (CTAB) protocol (Doyle and Doyle 1987). and resuspended in 100 µL TE. DNA amplification was performed using the ITS5/ITS4 primer set (White 1990), following the original PCR protocol with an annealing temperature of 56 °C, and by amplification of ∼1000 bp-fragment of the Large Subunit (LSU) of the nuclear ribosomal RNA gene, using the primers LR0R (5’-ACCCGCTGAACTTAAGC-3’) and LR5 (5’-TCCTGAGGGAAACTTCG-3’) following the original PCR protocol (Vilgalys and Hester 1990),using the DreamTaq DNA Polymerase (5 U/µL, Thermo Fisher Scientific, USA).

The amplified products were purified and sequenced by Eurofins Genomics (Ebersberg, Germany). Bacterial and fungal identification was performed using the highest identity score from BLAST 2.15.0 software of the National Center for Biotechnology Information (NCBI) against the NR database. Sequences were deposited in the GenBank database under accession numbers OQ874530, OQ874531, OQ874704, OQ874705, OQ883732-OQ883773 (Table 1). A percentage identity > 98% with NCBI sequences was accepted for species-level identification, while identities between 95% and 97% were classified at the genus level (Stackebrandt et al. 2022).

**Table 1.**
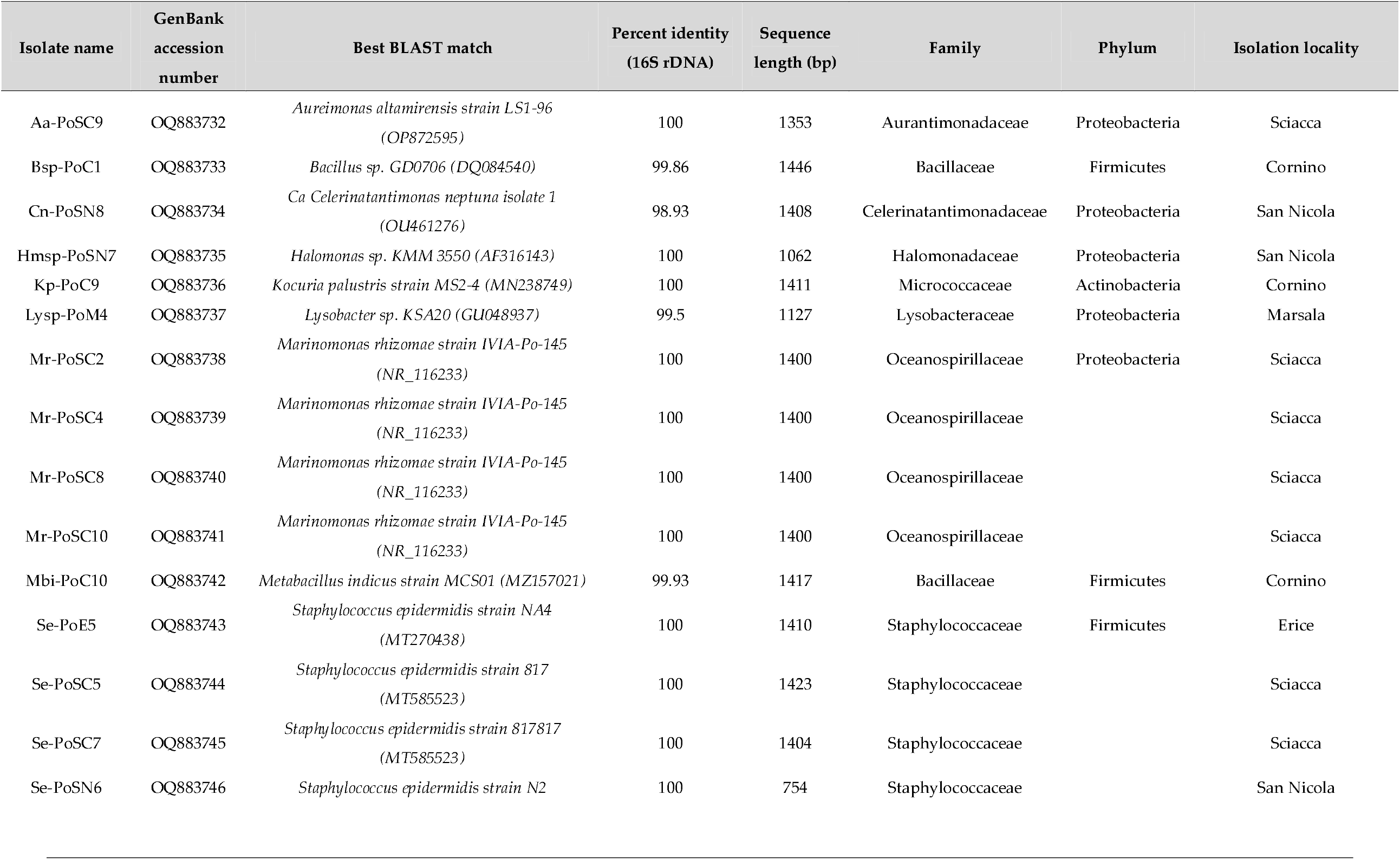
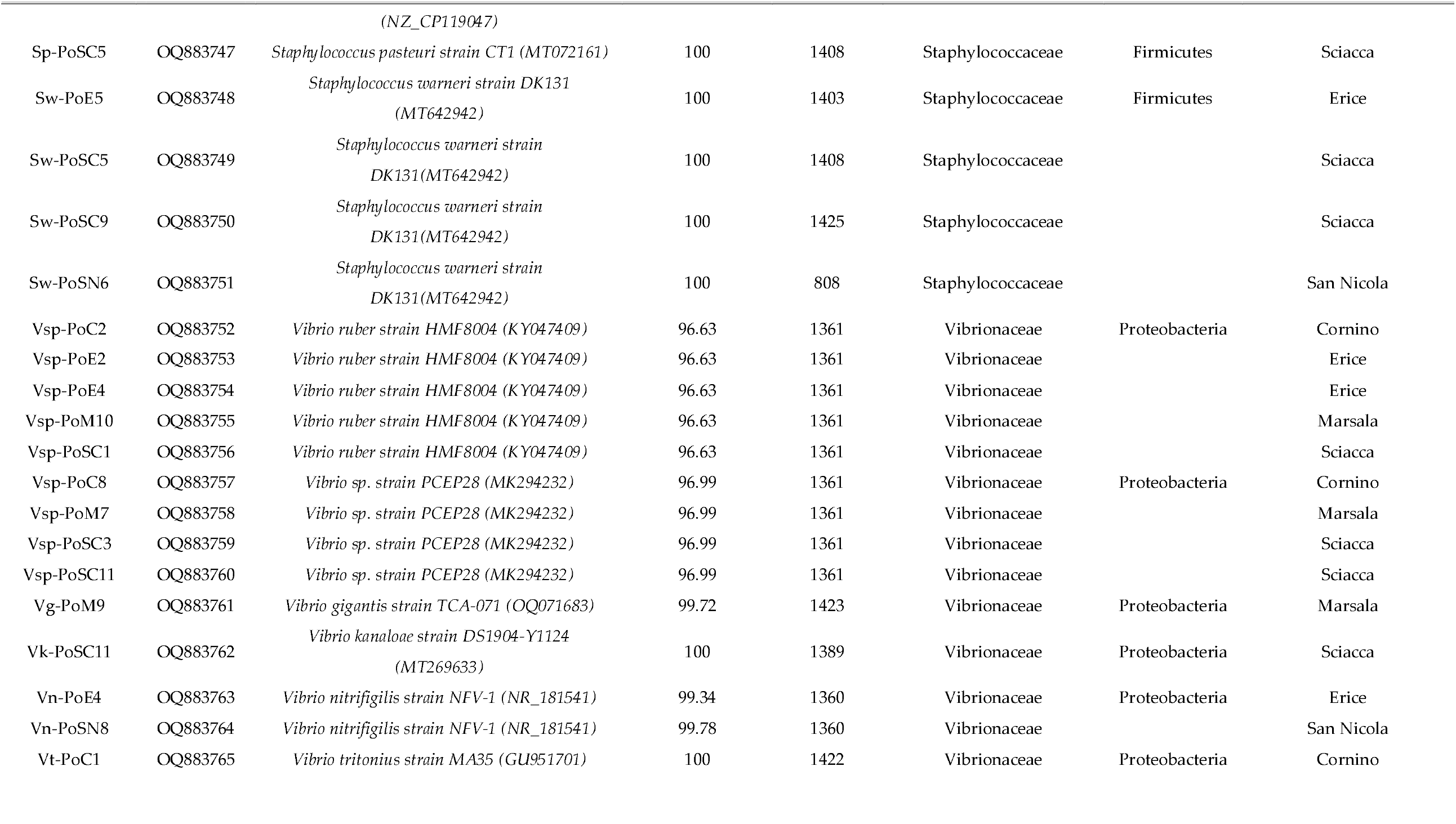
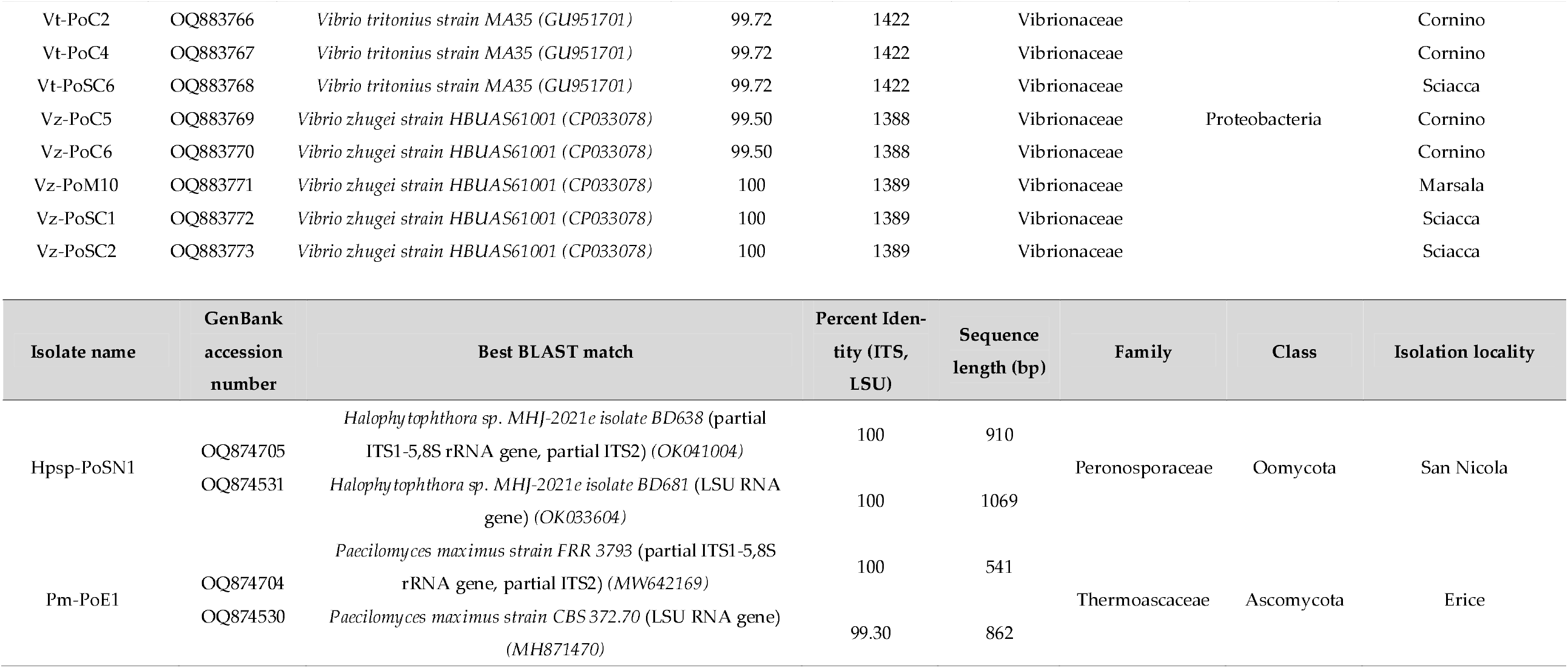
List of the 44 identified endophytes isolated from the *Posidonia oceanica* seeds with the corresponding molecular identification.

### 2.4. Determination of Plant Growth Promotion Properties and Enzymatic Activities of Bacterial Isolates

The production of indoleacetic acid (IAA) was evaluated using a colorimetric technique by incubating 3 mL of a liquid culture in Nutrient Broth (NB: 3 g yeast extract, 5 g peptone, 20 g NaCl, 200 mL ASW, SDW up to 1 L, pH 6.8-7.2) with L-tryptophan (100 mg/L), the IAA precursor, at 28 °C for 72 h in shaking condition at 120 rpm. Cultures were centrifuged for 5 min at 15,000 × g and supernatants transferred to glass tubes. Color changing to pink (positive reaction) was verified following the addition of 4 mL of Salkowski’s reagent (Gordon and Weber 1951). The amount of IAA was calculated by measuring the absorbance at 535 nm in a spectrophotometer (ASYS UVM-340 Microplate Reader, Biochrom, Ltd., Cambridge, UK), according to a standard curve appropriately realized.

Siderophores production was evaluated in Chrome Azurol S (CAS) agar plates (Schwyn and Neilands 1987). Aliquots (100 µL) of bacterial cultures were inoculated in the wells of the CAS plates and incubated for 7 days at 28 °C in darkness. Bacteria that produced siderophores showed an orange halo.

Phosphate solubilization was determined by the presence of a transparent halo around the bacterial cultures (100 µL) inoculated in wells of NBRIP plates (phosphate growth medium from the National Institute of Botanical Research) after 7 days at 28 °C (Nautiyal 1999).

Nitrogen fixation was assessed by plating strains in a nitrogen-free minimum medium (NFB; (Döebereiner 1995)) and incubating for 5 days at 28 °C. Bacterial growth indicated the possible ability of the bacteria to fix atmospheric nitrogen.

The formation of biofilms was determined by checking the adhesion capacity of the bacteria in microplates with 12 wells in NB at 28 °C for 4 days. After incubation, the biofilm formation was observed in the surface and/or bottom of the wells. Then, each well was stained for 20 min with 0.01% crystal violet (del Castillo et al. 2012) to evaluate the presence of a ring of biofilm around well walls.

The ACC-deaminase activity was performed as described by Penrose and Glick (2003) (Penrose and Glick 2003). Briefly, bacterial cultures were incubated in a salts minimal broth added with 5 mM 1-aminocyclopropane-1-carboxylic acid (ACC) for 24 h at 28 °C in shaking condition at 120 rpm. The α-ketobutyric acid produced was quantified using a standard curve with known concentrations by measuring absorption at 540 nm in a spectrophotometer. ACC-deaminase activity was expressed as µmoles of α-ketobutyrate per mg of protein per hour.

Enzymatic activities were determined on plates incubated at 28 °C for 7 days, observing the formation of halo around the bacterial biomass. DNAse activity was determined by streaking the isolates on DNAse agar plates (Scharlab, Spain) revealed with 1 M HCl. Amylase activity was performed on starch agar plates (Scharlab, Barcelona, Spain) and revealed with 10 mL lugol (Panreac Applichem, Spain). Protease and lipase activities were assessed in casein agar and Tween 80 media, respectively, as described by Harley and Prescott (2002) (Harley and Prescott 2002). Pectinase and cellulase activities were examined as described by Elbeltagy *et al*. (Elbeltagy et al. 2000). For pectinase activity, strains plated on ammonium mineral agar plates were revealed with 2% CTAB, and positive bacteria showed a halo. For cellulase activity, strains were plated on solid minimal medium supplemented with 0.2% yeast extract and 1% carboxymethylcellulose. Plates were developed by covering the plate with 1 mg/mL Congo Red solution (Sigma-Aldrich, USA) for 15 min and decolorizing with 1M NaCl for 20 min. Finally, chitinase activity was performed as described by Mesa *et al*. (Mesa et al. 2015) on agar plates containing minimal medium supplemented with colloidal chitin.

### 2.5. Data analysis

A non-parametric approach was adopted to analyse the culturable endophyte occurrence in seed material on NA and SGY media. To investigate the differences in CFU/g, a binomial sign non-parametric test was adopted (Higgins 2004) and implemented in Microsoft Excel (Microsoft Office 365).

To obtain information about the functional diversity of fungal strains, the FUNGuild database was used to estimate the trophic modes and guilds of fungi (Nguyen et al. 2016). This tool allows to predict the following primary fungal lifestyles: pathotroph: receiving nutrients at the expense of the host cells and causing disease; saprotroph: receiving nutrients by breaking down dead host cells; symbiotroph: receiving nutrients by exchanging resources with host cells.

The Venn diagrams were constructed manually or using the webtool: https://bioinformatics.psb.ugent.be/webtools/Venn/.

## 3. Results

### 3.1. Endophyte isolation

The number of bacterial and fungal colonies on NA and SGY media increased over the incubation period, with most colonies appearing within the first 10 days. No colonies were observed on any of the negative control plates. Similarly, no growth was obtained from some seed samples regardless of collection site and media (Figure 2). The absence of colonies on both NA and SGY plates was most likely due to the low microbial concentrations typically associated with plant endophytes. Colony-forming units were observed from 30 seed samples on SGY plates (68% of seeds) and 21 seed samples on NA plates (48%) (Figure 2, Appendix 1). CFUs were recovered on both media from 19 seed samples (43%), exclusively from SGY in 11 samples (25%), and exclusively from NA in 2 samples (5%) (Figure 2, Appendix 1). Twelve out of 44 samples (27%) showed no colony growth on either medium (Figure 2).

**Figure 2.**
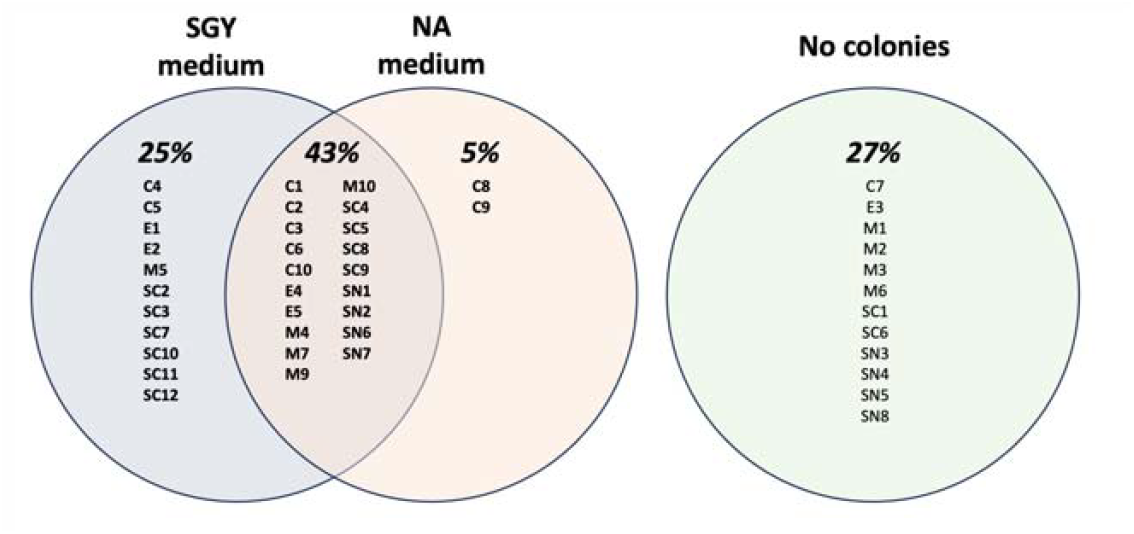
Percentage and identity of seeds yielding colonies on SGY and NA media, or none.

The highest total endophyte concentrations, expressed as CFU/g, were 3.62 × 10^2^ on SGY and 3.39 × 10^2^ on NA (Appendix 1). Statistical analysis revealed no significant difference in total endophyte concentrations between the two media at a confidence level (1–α) ≥ 0.95.

### 3.2. Molecular identification of culturable endophytes

Based on phenotypic characteristics, 42 bacterial and two fungal isolates were selected (Appendix 2) and identified through sequence analysis. Bacterial 16S rDNA sequences ranged from 754 to 1446 bp in length, while fungal ITS and LSU regions ranged from 541 to 910 bp and 862 to 1069 bp, respectively (Table 1). The identified endophytic bacterial isolates belonged to three phyla—Proteobacteria, Firmicutes, and Actinobacteria—and were associated with ten different genera (Table 1). Most isolates were grouped into three dominant genera: *Vibrio, Staphylococcus*, and *Marinomonas*, followed by *Aureimonas, Bacillus, Celerinatantimonas, Halomonas, Kocuria, Lysobacter*, and *Metabacillus* (Figure 3).

**Figure 3.**
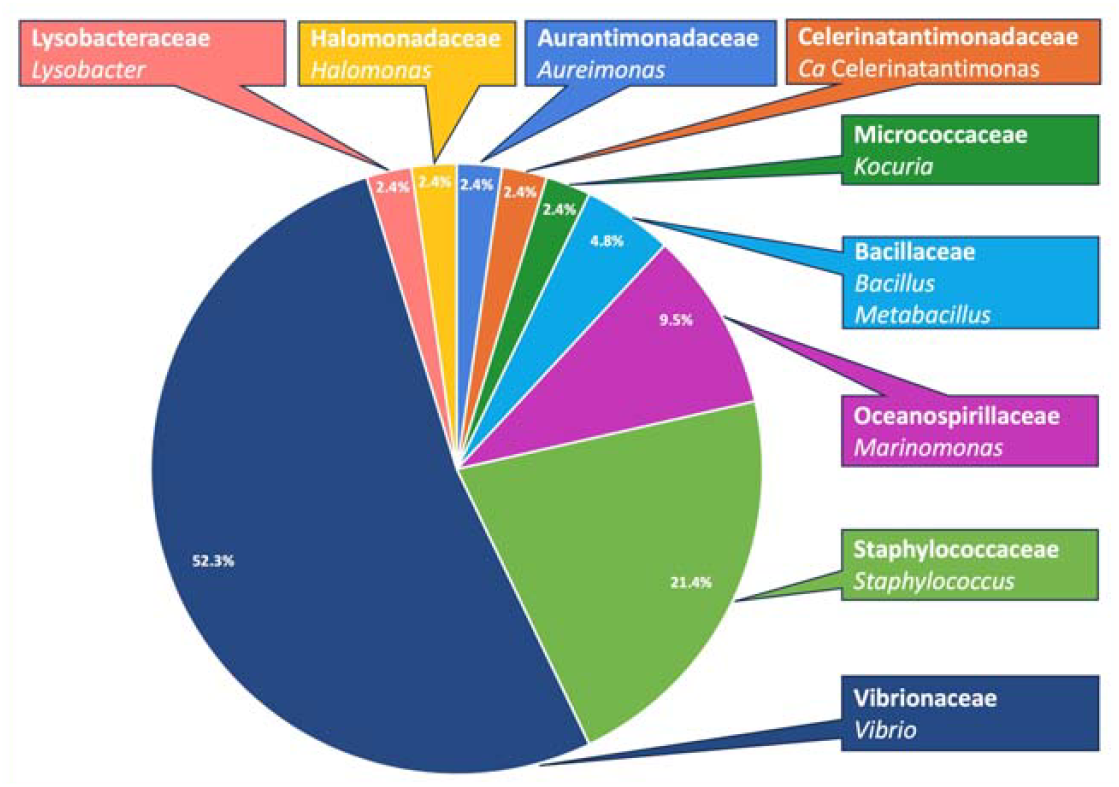
Distribution of bacterial families isolated from seeds.

In terms of sequence identity, 78.6% of the isolates showed high similarity (≥98 %) to 16S rDNA sequences available in GenBank (Table 1). Nine *Vibrio* isolates, however, exhibited lower identity values (<97%) with reference sequences in GenBank (Table 1). Specifically, five isolates (Vsp-PoC2, Vsp-PoE2, Vsp-PoE4, Vsp-PoM10, and Vsp-PoSC1) were identical and associated with *Vibrio ruber* strain HMF8004 (GenBank accession KY047409, 96.63%). The remaining four *Vibrio* isolates (Vsp-PoC8, Vsp-PoM7, Vsp-PoSC3, Vsp-PoSC11) were identical and associated with *Vibrio* sp. strain PCEP28 (GenBank MK294232, 96.99%).These two groups of *Vibrio* isolates shared 95.96% identity with each other.

In total, two fungal colonies with distinct growth rates and morphological characteristics were isolated from two seed samples collected in San Nicola and Erice, respectively (Appendix 2). The isolates Pm-PoE1 and Hpsp-PoSN1 showed 100% identity with fungal ITS sequences deposited in GenBank, corresponding to *Paecilomyces maximus* strain FRR 3793 (GenBank MW642169) and *Halophytophthora* sp. MHJ-2021e isolate BD638 (GenBank OK041004), respectively (Table 1). LSU sequence analysis further confirmed species identification, showing 99.3% identity between Pm-PoE1 and *P. maximus* strain CBS372.70 (GenBank MH8714670), and 100% identity between Hpsp-PoSN1 and *Halophytophthora* sp. MHJ-2021e isolate BD638 (GenBank OK033601).

### 3.3. PGP properties: Plant Growth Promoting Traits and enzymatic activities

A total of 22 bacterial isolates (Figure A1)—selected based on their higher in vitro growth capacity—were characterized to assess the presence of plant growth-promoting (PGP) traits and enzymatic activities (Tables 2 and 3).

**Table 2.** PGP properties showed by the endophytes isolated from *Posidonia oceanica* seeds.

| Isolate | Identification | IAA pro-<br>duction | Siderophore<br>production | Phosphate<br>solubilisation | Nitrogen<br>fixation | Biofilm |  |  | ACC deaminase<br>activity |
| --- | --- | --- | --- | --- | --- | --- | --- | --- | --- |
|  |  |  |  |  |  | Surface | Bottom | Ring |  |
| Aa-PoSC9 | *Aureimonas altamirensis* | - | 0.4 | 0.4 | + | - | +++ | - | - |
| Bsp-PoC1 | *Bacillus sp.* | - | 0.3 | 0.2 | - | + | ++ | - | - |
| Hmsp-PoSN7 | *Halomonas sp.* | 4.61 | 2.5 | - | - | - | ++ | ++ | - |
| Kp-PoC9 | *Kocuria palustris* | - | 0.1 | - | - | - | - | - | - |
| Lysp-PoM4 | *Lysobacter sp.* | - | 0.3 | - | - | - | ++ | - | - |
| Mr-PoSC8 | *Marinomonas rhizomae* | 3.39 | - | - | - | - | + | - | - |
| Mbi-PoC10 | *Metabacillus indicus* | 0.51 | - | - | + | - | ++ | - | - |
| Vsp-PoC2 | *Vibrio sp.* | 0.98 | - | - | - | ++ | ++ | ++ | - |
| Vsp-PoE2 | *Vibrio sp.* | - | 0.8 | - | + | + | - | - | - |
| Vsp-PoE4 | *Vibrio sp.* | - | 0.9 | - | + | +++ | - | - | - |
| Vsp-PoM10 | *Vibrio sp.* | 0.51 | 0.1 | - | - | - | + | ++ | - |
| Vsp-PoM7 | *Vibrio sp.* | - | 0.8 | - | + | - | +++ | + | - |
| Vsp-PoSC11 | *Vibrio sp.* | - | 0.9 | - | + | +++ | + | - | - |
| Vg-PoM9 | *Vibrio gigantis* | - | 0.2 | - | - | - | + | - | - |
| Vk-PoSC11 | *Vibrio kanaloae* | - | - | - | - | - | + | - | - |
| Vn-PoE4 | *Vibrio nitrifigilis* | - | 0.5 | 0.1 | - | - | + | - | - |
| Vn-PoSN8 | *Vibrio nitrifigilis* | 5.9 | 0.7 | 0.6 | - | - | + | - | - |
| Vt-PoC4 | *Vibrio tritonius* | - | - | - | - | - | - | - | - |
| Vt-PoSC6 | *Vibrio tritonius* | - | 0.2 | - | - | - | + | - | - |
| Vz-PoC5 | *Vibrio zhugei* | 0.69 | 0.3 | - | - | - | - | - | - |
| Vz-PoM10 | *Vibrio zhugei* | - | 0.4 | - | - | + | - | + | - |
| Vz-PoSC1 | *Vibrio zhugei* | 1.77 | 0.1 | - | - | - | +++ | + | - |
|  |  |  |  |  |  | Surface | Bottom | Ring |  |
| Aa-PoSC9 | *Aureimonas altamirensis* | - | 0.4 | 0.4 | + | - | +++ | - | - |

**Table 3.**
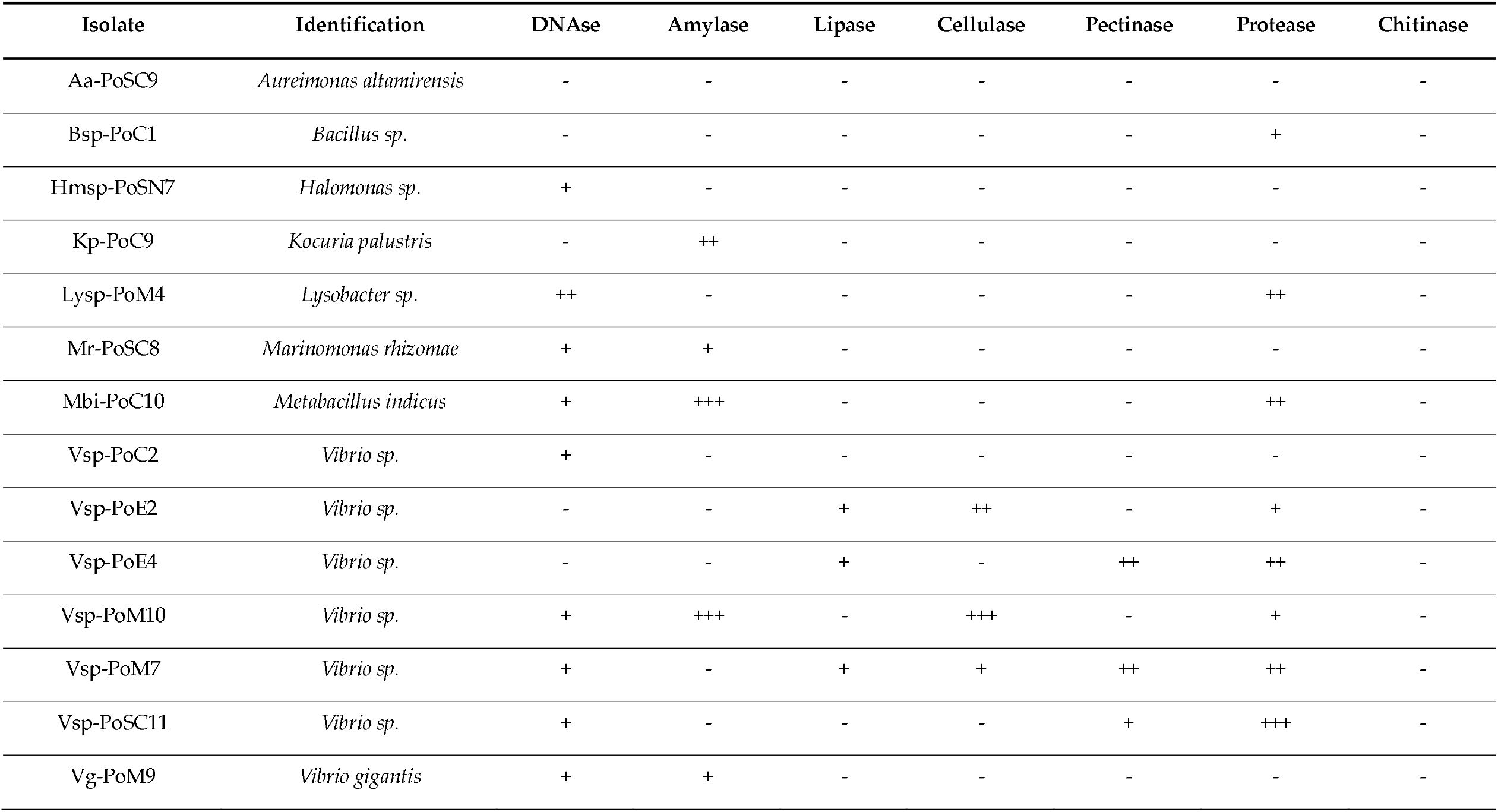
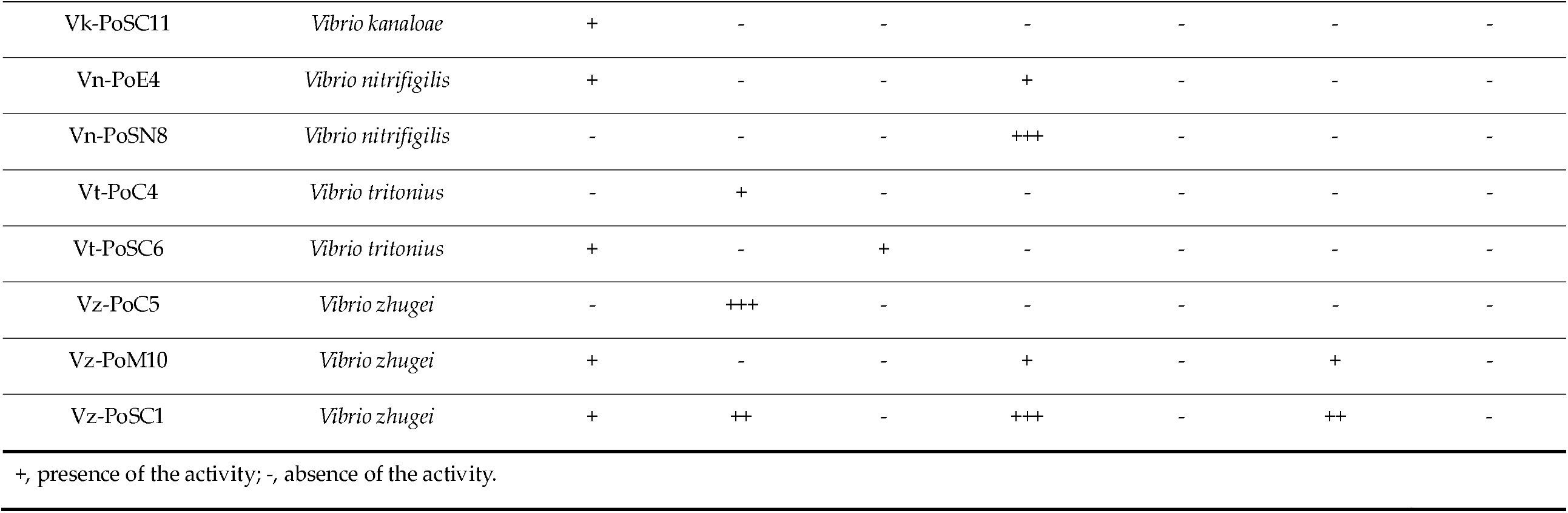
Enzymatic activities showed by the endophytes isolated from *Posidonia oceanica* seeds.

All isolates exhibited at least one of the tested properties Most isolates (77%) displayed between four and eight traits simultaneously. Only one isolate (Vt-PoC4) did not show any PGP trait, and one (Aa-PoSC9) no enzymatic activity.

The most frequently observed PGP characteristics were biofilm formation (86%), siderophore production (77%), and indole-3-acetic acid (IAA) production (36%), followed by nitrogen fixation (22%), and phosphate solubilization (18%) (Figure 4A). Most isolates formed biofilms predominantly at the bottom of the wells (Table 2). Isolate Hmsp-PoSN7 (*Halomonas* sp.) exhibited the highest siderophore production, as indicated by the halo diameter on CAS medium, while isolate Vn-PoSN8 (*Vibrio nitrifigilis*) was the most efficient IAA producer. None of the bacterial isolates exhibited ACC deaminase activity under in vitro conditions.

**Figure 4.**
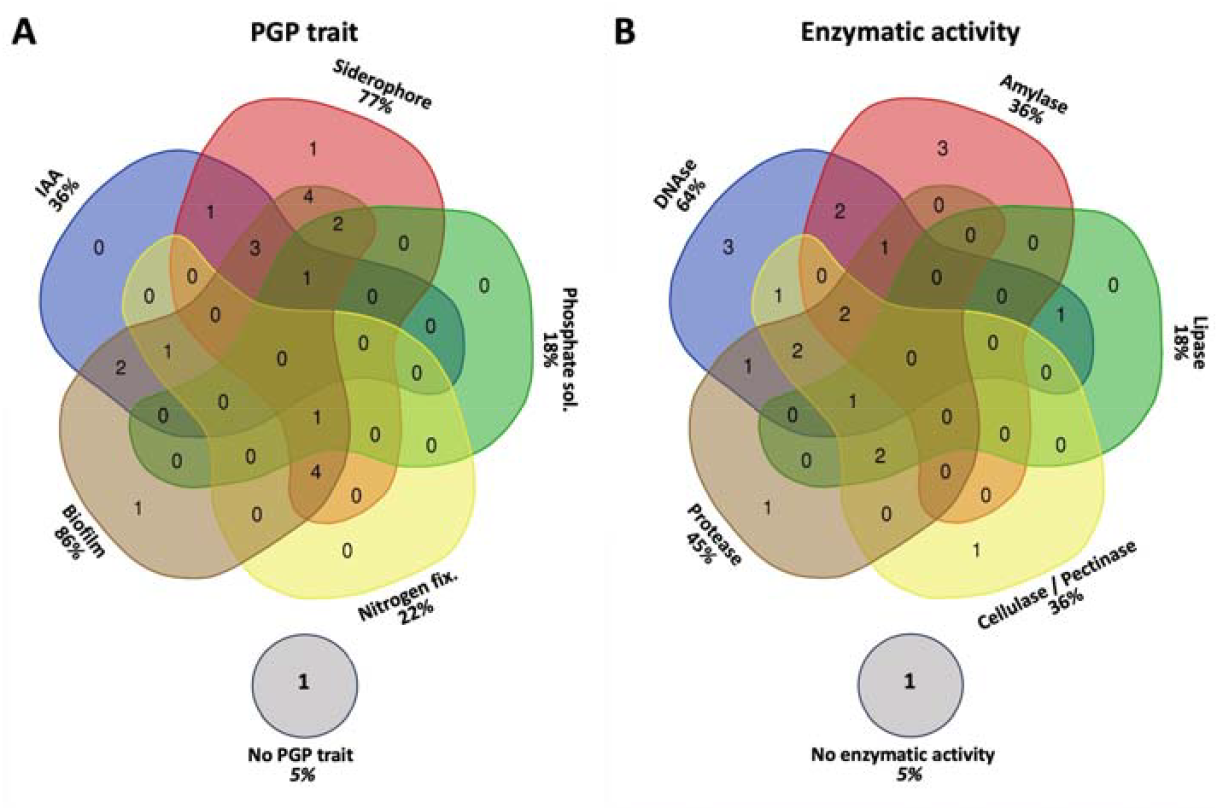
Frequencies of PGP traits (A) and enzymatic (B) activities among the 22 bacterial isolates. Cellulase and pectinase activities were grouped together.

Among the six enzymatic activities tested, DNase (64%) was the most prevalent, followed by protease (45%), amylase (36%), cellulase (32%), lipase (18%) and pectinase (13%) (Figure 4B and Table 3). The strongest amylase and cellulase activities were both observed in isolate Vsp-PoM10. None of the bacterial isolates exhibited chitinase activity under in vitro conditions.

### 3.4. Trophic mode of endophytic fungi

Fungal taxonomic and functional analyses using FUNGuild classified the two fungal isolates, Pm-PoE1 and Hpsp-PoSN1, into distinct putative trophic modes. *P. maximus* was assigned to the pathotroph–saprotroph–symbiotroph mode, whereas *Halophytophthora* sp. was categorized as a saprotroph. The former trophic mode is primarily associated with endophytes, clavicipitaceous fungi, plant pathogens, and both plant and wood saprotrophs; while the latter is mainly represented by plant and wood saprotrophs.

## 4. Discussion

The positive effects of plant–microorganism interactions, widely demonstrated in terrestrial agroecosystems (Compant et al. 2019; Pacifico et al. 2019), are now increasingly being recognized in marine environments, supported by the growing body of knowledge on the seagrass holobiont (Ugarelli et al. 2017; Tarquinio et al. 2019). Early experiments with epiphytic microorganisms associated with the vegetative organs of *P. oceanica* revealed their ability to stimulate plant development and influence the structure of the epiphyte community (Celdrán et al. 2012). More recently, positive effects on plant development were also ascribed to endophytes (Mohr et al. 2021). Growing evidence indicates plant-associated microbes as key factors enhancing development and stress tolerance in *P. oceanica* meadows. However, it is not known how these beneficial guests are transmitted among individual plants. While rhizomes can extend laterally and interconnect plants within the same meadow, it is still an open question whether different meadows are characterized by similar epiand endophytes, and how these are transmitted through generations. In terrestrial plants, it is known that seeds can act as a trans-generational reservoir of endophytes (Truyens et al. 2015). In *P. oceanica*, however, no study had ever described the presence of endophytes. In our work, we present the diverse culturable bacterial and fungal endophytic community isolated from *P. oceanica* seeds collected in different Sicilian locations. Located in the centre of the Mediterranean, at the intersection of the western and Levantine gene pools, Sicilian meadows present the highest genetic biodiversity (Arnaud-Haond et al. 2007). By sampling seeds along the different quadrants of the Sicilian coasts, we aimed to further maximise the potential recovery of microbial endophyte communities. Since the seeds were collected from stranded fruits, it was not possible to trace each seed back to a specific mother plant or meadow. However, it is likely that the fruits had detached from nearby meadows. Whether a mother plant consistently transmits the same endophytic community to all its seeds remains an open question.

Our primary objective was to isolate culturable endophytic microorganisms that could enhance the growth and stress tolerance of seagrass propagation material, with the aim of facilitating restoration efforts. Due to the lack of standardized protocols for isolating endophytes from *P. oceanica* seeds, we used two culture media recommended in the literature for marine microbial isolation (Kiki 2016). Based on CFU/g values, both media proved effective for isolating seed-associated endophytes. Nonetheless, some samples yielded no CFUs on either medium, likely due to either a low microbial load or selective limitations of the media—particularly SGY, which has a more complex formulation (Kiki 2016). Isolation efficiency likely depends not only on culture conditions and disinfection protocols but also on the inherent rarity, uneven distribution, and varied growth capacities of endophytes under aseptic conditions (Rungjindamai and Jones 2024). Future isolation efforts could benefit from the use of selective, enriched, and oligotrophic media to increase recovery rates and capture slow-growing taxa.

A significant number of seeds did not yield colonies on plates. However, this does not necessarily indicate seed sterility, as our study focused exclusively on the culturable fraction of the microbiota. Furthermore, we only tested two generalist media, which may not support the growth of more specialized microbial species. An additional level of complexity might derive from the competition among species in the same plate. Although we streaked colonies in fresh plates multiple times to isolate pure species, we cannot exclude that in the original homogenate, the presence of some microbes inhibited the growth of others. Therefore, further complementary metagenomic approaches will be necessary to detect the presence of unculturable or elusive microorganisms.

The bacterial isolates identified in *P. oceanica* seeds largely belong to the same phyla reported in seeds of the seagrass *Halophila ovalis* (Tarquinio et al. 2021), suggesting that unrelated seagrass species—belonging to lineages that diverged approximately 30 million years ago and independently adapted to marine environments (Lee et al. 2018) may host similar endophytic communities. Notably, our study is the first to report the presence of both Oomycota and Ascomycota in seagrass seeds. Our PGP assay results suggest that *P. oceanica* seeds, in addition to containing a large amount of reserves within the endosperm, also recruit and vertically transmit microorganisms capable of supporting germination and early seedling development, through hormone production or by facilitating the acquisition of essential nutrients such as iron and nitrogen.

Among the bacterial genera identified in this study, *Marinomonas, Celerinatantimonas*, and *Vibrio* are commonly found in aquatic and marine environments. *Marinomonas* has previously been reported as an endophyte in *P. oceanica*, and several species, including *M. rhizomae*, are known to be abundant within *P. oceanica* tissues (Espinosa et al. 2010; Lucas-Elió et al. 2011; Garcias-Bonet et al. 2012). Growth-promoting *Marinomonas* species have also been detected in the core microbiome of the seagrass *Halophila ovalis* seeds (Tarquinio et al. 2021). Given the reported positive effects of *Marinomonas* on seedling growth (Celdrán et al. 2012) and the strong IAA production observed in our *M. rhizomae* isolate Mr-PoSC8, our findings support the hypothesis of a close plant–microbe association that may be sustained through vertical transmission.

Recently, Mohr *et al*. (Mohr et al. 2021) demonstrated the positive outcomes of the symbiotic association between *P. oceanica* and the nitrogen-fixing endophyte *Candidatus* Celerinatantimonas neptuna, which colonizes the plant roots. Other nitrogen-fixing species, including *C. yamalensis* and *C. diazotrophica*, have been isolated from the seagrass *Thalassia hemprichii* and from estuarine grasses such as *Spartina alterniflora* and *Juncus roemerianus*, respectively (Cramer et al. 2011; Mohr et al. 2021). As with terrestrial plants, these symbiotic relationships enable marine species to thrive in nitrogen-poor, oligotrophic environments, such as seagrass meadows and salt marshes. Our isolate Cn-PoSN8 showed 98.93% nucleotide identity with a ∼1400 bp fragment of the *Candidatus* C. neptuna genome (GenBank OU461276.1, region 426558–425198) retrieved from *P. oceanica* roots in the Elba/Pianosa area. We only found *Celerinatantimonas* in one locality, San Nicola, located on the southern Tyrrhenian Sea, while it was absent in seeds collected in west Sicily and along the Strait of Sicily. *C. neptuna* was originally found in roots of *P. oceanica* meadows from the Tuscan archipelago, in northern Tyrrhenian. It is therefore intriguing to question whether this important endosymbiont is restricted to meadows of the Tyrrhenian Sea, possibly to the western genetic pool of *P. oceanica*, or its presence is widespread in the whole basin. Its presence in *P. oceanica* seeds suggests a potential route of vertical transmission, likely reflecting its beneficial role for the host (Mohr et al. 2021). Although we were unable to evaluate the PGP traits of this isolate due to its failure to survive long-term storage at –80 °C—possibly because it may require mesophilic and partially anaerobic growth conditions (Cramer et al. 2011)—we successfully identified other nitrogen-fixing bacteria in seeds collected from different Sicilian sites: *Aureimonas, Metabacillus*, and different *Vibrio* isolates obtained in this study displayed nitrogen-fixing capacity.

In this study, we identified five *Vibrio* sequence groups corresponding to known species not previously reported as seagrass endophytes, and nine isolates grouped into two sequence groups with <97% identity to *Vibrio* 16S rDNA sequences available in the GenBank. Further multilocus sequence typing (MLST) will be necessary to determine the precise taxonomic status of these isolates, which may represent two novel species (Jiang et al. 2022). Numerous studies have reported associations between *Vibrio* spp. and marine micro- and macroalgae (Sampaio et al. 2022). Some *Vibrio* species are also known to colonize seagrass canopies (e.g., *Zostera* spp.), contributing to the reduction of microbial loads in surrounding water bodies (Reusch et al. 2021). Additionally, endophytic *Vibrio* spp. have been isolated from the mercury-tolerant halophyte *Halimione portulacoides* (Fidalgo et al. 2016). A synergistic combination of *Vibrio aestuarianus* (a nitrogen-fixing bacterium) and *Vibrio proteolyticus* (a phosphate-solubilizing species), both present in the mangrove rhizosphere, significantly enhanced growth in the halophyte *Salicornia* (Bashan et al. 2000). Although knowledge about the role of *Vibrio* spp. as plant endophytes remains limited, several species identified in this study have already been associated with plant-beneficial traits, particularly those linked to nutrient acquisition (Huang et al. 2021). Here, we confirmed the presence of multiple PGP properties in nearly all *Vibrio* isolates obtained from *P. oceanica* seeds, with *V. europaeus* and *V. zhugei* standing out as the most promising candidates.

Most of the remaining culturable bacteria isolated from *P. oceanica* seeds have previously been identified in plant species adapted to extreme environments, such as saline or contaminated soils. *Halomonas* sp., *Staphylococcus warneri, S. epidermidis*, and *Kocuria palustris* have been reported as members of the endophytic microbiota of the halophyte *Seidlitzia rosmarinus*, which grows in saline soils (Shurigin et al. 2020).

Halotolerant strains of *Metabacillus indicus* and *S. warneri* exhibiting multiple PGP traits were also isolated from the rhizosphere of the halophyte *Sesuvium portulacastrum* (John et al. 2023). *Halomonas* spp. are frequently recovered from roots, leaves (Zhang et al. 2018; Kearl et al. 2019), and seeds of halophytic species (Wang et al. 2021). Their ability to adapt to harsh environmental conditions is largely attributed to specific genomic features (Chaudhry et al. 2017; Ali et al. 2022). For instance, Zhang *et al*. (Zhang et al. 2020) showed that endophytic *Halomonas* strains respond to salt stress by modulating over 800 genes involved in cellular and metabolic processes. Our results are fully consistent with the literature, confirming the presence of beneficial traits in *Halomonas* and *Kocuria* isolates obtained from seeds of a marine phanerogam. The *Kocuria* isolate Kp-PoC9 identified in this study belongs to a group of endophytes characterized by a limited number of known beneficial traits. However, we do not exclude that its recruitment into the seed microbiome may be linked to other advantageous properties. In fact, some *Kocuria* species have been identified as arsenic-resistant endophytes, potentially supporting host plants in environments contaminated with arsenic and other heavy metals (Román-Ponce et al. 2016; Zacaria Vital et al. 2019). *K. palustris* isolated from *P. oceanica* exhibited high sequence identity with the arsenic-resistant NE1RL3 strain from *Sphaeralcea angustifolia* growing in metal-contaminated soils (Zacaria Vital et al. 2019).It is possible that its presence in seagrass seeds could assist seedlings in colonizing polluted sites. Furthermore, this species may offer additional benefits through the production of bioactive compounds with fungicidal activity, as reported for *K. palustris* strains isolated from marine sponges effective against *Fusarium oxysporum* (Setiawan et al. 2022).

Although *Staphylococcus* spp. are primarily known as potential human and animal pathogens, they have also been frequently isolated from vegetative organs (Surette et al. 2003; Chiellini et al. 2015; Gabriele et al. 2022) and seeds (Liu et al. 2013; Faddetta et al. 2021) of various plants. According to the literature, *Staphylococcus* strains adapted to an endophytic lifestyle differ from human strains due to the presence of gene clusters involved in the DNA damage caused by ROS accumulation and UV radiation (Chaudhry and Patil 2016). In the case of our isolates, due to the lack of complete genomic data and information on the presence of virulence factors, we decided to exclude them from PGP trait analyses, as no immediate biotechnological application can be foreseen for *P. oceanica* or agricultural crops.

*Bacillus* and *Metabacillus* include many well-known plant endophytes (Zhang et al. 2018; Yin et al. 2022). Many *Bacillus* spp. are already used in commercial bioformulations due to their broad plant-beneficial effects, including protection against pathogens and stimulation of plant growth (Lopes et al. 2018). In addition to these genera, the presence of *Lysobacter* in *P. oceanica* seeds is particularly noteworthy due to its potential as a biological control agent (BCA). *Lysobacter* spp. are part of complex microbial communities associated with soil and plants and are capable of colonizing a wide range of extreme environments, including aquatic systems ((Brescia et al. 2020). Some endophytic *Lysobacter* strains have already demonstrated BCA activity against several plant pathogens through the production of lytic enzymes and antibiotic compounds (Drenker et al. 2023; Tu et al. 2023). Our study confirms the presence of these three genera as endophytes in a marine plant and places them among the isolates with the highest number of PGP properties.

Few *Aureimonas* species have been reported as plant endophytes. *A. endophytica* was isolated in China from the halophyte *Anabasis elatior* (Liu et al. 2016), while *A. altamirensis* was recovered from leaves of ash trees tolerant to dieback caused by *Hymenoscyphus fraxineus* (Becker et al. 2022). Combining functional genome analysis and inoculation assays, Becker *et al*. (Becker et al. 2022) demonstrated that this species may contribute to increased plant resilience. Our findings provide additional evidence for the presence of *Aureimonas* as a plant endophyte with beneficial properties, including nitrogen-fixing capacity. However, we cannot exclude a potential antagonistic role, which will require further investigation through specific bioassays.

Recent studies have reported several endophytic fungal species associated with the vegetative organs of *P. oceanica* in the central Mediterranean (Torta et al. 2015, Torta et al. 2022; Vohník et al. 2016, 2019). However, only a limited number of culturable fungal and oomycote species were recovered from seeds in the present study. The two genera identified had previously been isolated from surface-sterilized *P. oceanica* tissues (Torta et al. 2022).

Fungal genera most frequently detected as endophytes in roots and rhizomes—such as *Posidoniomyces, Lulwoana, Ochroconis, Penicillium*, and members of the Xylariaceae family (Torta et al. 2015, Torta et al. 2022; Vohník et al. 2019; Vohník 2022)—were not isolated from seeds. According to Torta *et al*. (Torta et al. 2022), fungal endophyte distribution varies among plant organs in *P. oceanica*, with higher colonization in basal structures (roots and rhizomes) and only sporadic presence in leaves. Based on our findings, we cannot exclude that fungal endophytes are naturally scarce in seeds or that they exhibit a species composition distinct from other plant organs, particularly the basal ones. Given the important role of root-associated fungal endophytes in maintaining plant health (Torta et al. 2022), further research is needed to determine the timing and mode of root colonization.

*Paecilomyces* species are found in a wide range of terrestrial environments. While many are known as biological control agents against pests and pathogens, others have also been described as PGP fungi (Moreno-Gavíra et al. 2020). Some strains of *P. maximus* (syn. *P. formosus*; (Houbraken et al. 2020) have been linked to tree declines in both natural and cultivated forests (Sabernasab et al. 2019; Ozan et al. 2022). This species is currently considered a complex comprising at least three cryptic taxa (Houbraken et al. 2020), potentially showing a dual role in plants—acting as beneficial endophytes by promoting growth through hormone production (IAA and GA) and increasing tolerance to abiotic stresses, especially heavy metals (Bilal et al. 2017). Therefore, the presence of *P. maximus* in *P. oceanica* seeds may confer developmental advantages and reduce the negative effects of heavy metal contamination.

The genus *Halophytophthora* includes species reported as saprotrophs or pathogens in mangrove forests, lagoons, estuaries, salt marshes, and marine ecosystems (Maia et al. 2022). *Halophytophthora* sp. Zostera infection in the seagrass *Zostera marina* has been associated with reduced seed viability and seedling growth, posing a threat to coastal ecosystem restoration (Govers et al. 2016). Recently, three *Halophytophthora* species have been isolated in rotting *P. oceanica* seeds and postulated to be the causative agents of early seed mortality in this seagrass (Alagna et al. 2024). When cast seeds are collected and placed in aquaria, a fraction of these, about 10-45%, cover in whitish mould and rot. Treatment with antifungal chemicals was tested as a mean to control infection (Alagna et al. 2024). Our findings indicate that this saprotroph manages to spread, spatially and through generations, as a seed endophyte, and that protocols of seed disinfection are likely not able to prevent infection by this fungus. Interestingly, since we found *Halophytophthora* only in few seeds, it is possible that the fraction of seeds that rot in the first weeks after collection is already carrying the fungus as endophyte, and is therefore “doomed”.

Clarifying the potential pathogenicity of *Halophytophthora* sp. and *P. maximus* in Mediterranean seagrasses will require additional research, including isolation from symptomatic plants and experimental inoculation of seeds and seedlings under controlled conditions. Furthermore, field studies combining culture-dependent and metagenomic approaches are needed to investigate the spread of fungal endophytes via *P. oceanica* seeds and to assess their role in the epidemiology of potential plant pathogens.

Among the long-term consequences of climate change, the decline in the availability and quality of water bodies resources represents one of the most critical challenges for global agriculture. Water is an indispensable component of food production; according to UN and FAO estimates, approximately 3,000-5,000 litres of water are required to satisfy the daily dietary needs of a single person. Furthermore, the Global Risks Report of the World Economic Forum identifies water crises as the third most severe global threat in terms of impact on humanity (Pathirana and Carimi 2024). In this scenario, salinity has emerged as one of the major abiotic constraints limiting crop productivity and quality. The Food and Agriculture Organization (FAO 2024) estimates that around 1.4 billion hectares - over 10% of the global land surface - are already affected by salinization, with an additional one billion hectares at imminent risk due to climate change and unsustainable management practices. Our findings highlight that endophytic microorganisms associated with *P. oceanica* represent a valuable and largely untapped source of plant-beneficial microbes. Cultivable endophytes isolated from *P. oceanica* display traits potentially linked to salinity tolerance, such as the production of plant growth–promoting substances and allow to guarantee nitrogen fixation even in situations of saline stress. The use of these endophytes as bioinoculants in terrestrial crops could therefore contribute to enhancing plant resilience under saline conditions, improving both productivity and sustainability in salt-affected soils.

## 5. Conclusions

This study unravels the mechanisms of transgenerational and spatial diffusion of endophytic communities in *P. oceanica* meadows. It also expands the list of culturable endophytes from this species, providing a thorough characterization of their putative beneficial services. Most of the isolated microbial species exhibit traits associated with plant nutrition and growth promotion. In addition, isolate selection revealed the co-occurrence of different microbial species within the same seed, potentially capable of performing complementary or overlapping functions. Their impact on seedling development and pathogen resistance will be further evaluated through in planta trials under controlled conditions, laying the groundwork for their potential use in strategies aimed at protecting and restoring this dominant Mediterranean seagrass. Additionally, the strains isolated and characterized in this work are fully culturable in vitro, allowing a scalable application as inoculant in other plant species, such as other seagrasses, but also terrestrial crops, potentially enhancing tolerance to stress such as saline soils.

## Author Contributions

Dalila Crucitti: Conceptualization (equal); methodology (lead); validation (lead); formal analysis (lead); investigation (lead); resources (supporting); data curation (lead); writing—original draft preparation (lead); writing—review and editing (lead); visualization (equal).

Alberto Sutera: Investigation (equal); resources (equal); visualization (supporting).

Roberto De Michele: Conceptualization (supporting); formal analysis (equal); resources (lead); writing—original draft preparation (equal); writing—review and editing (equal); visualization (lead); project administration (supporting); funding acquisition (lead).

Francesco Carimi: Resources (supporting); writing—review and editing (equal); visualization (equal); project administration (lead); funding acquisition (equal).

Stefano Barone: Formal analysis (supporting); data curation (supporting).

Fabio Badalamenti: Resources (equal); review and editing (supporting); project administration (equal).

Davide Pacifico: Conceptualization (lead); methodology (equal); validation (equal); data curation (equal); writing—original draft preparation (equal); writing—review and editing (equal); visualization (equal).

## Acknowledgements

This research was funded by the project Marine Hazard, PON03PE_00203_1 (Ministry of Education, University and Research, MUR, Italy

## Data availability

The datasets generated and analyzed during the current study are available in the GenBank repository with the accessions listed in Table1.

## Ethics statement

None required

## Appendices

### Appendix 1 Weight of *Posidonia oceanica* seeds, total colonies counted and Colony Forming Unit (CFU)/g of fresh material on SGY and NA media

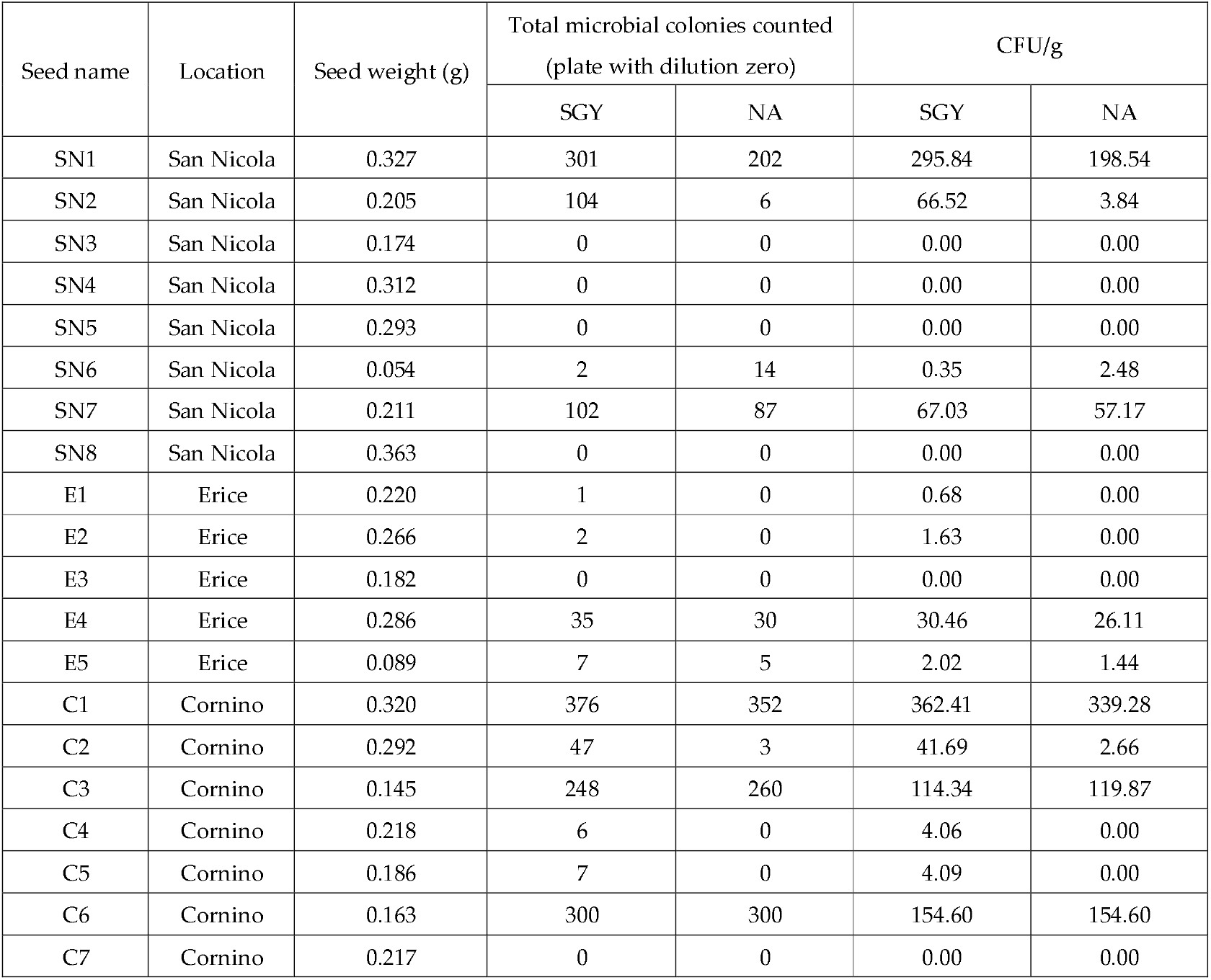
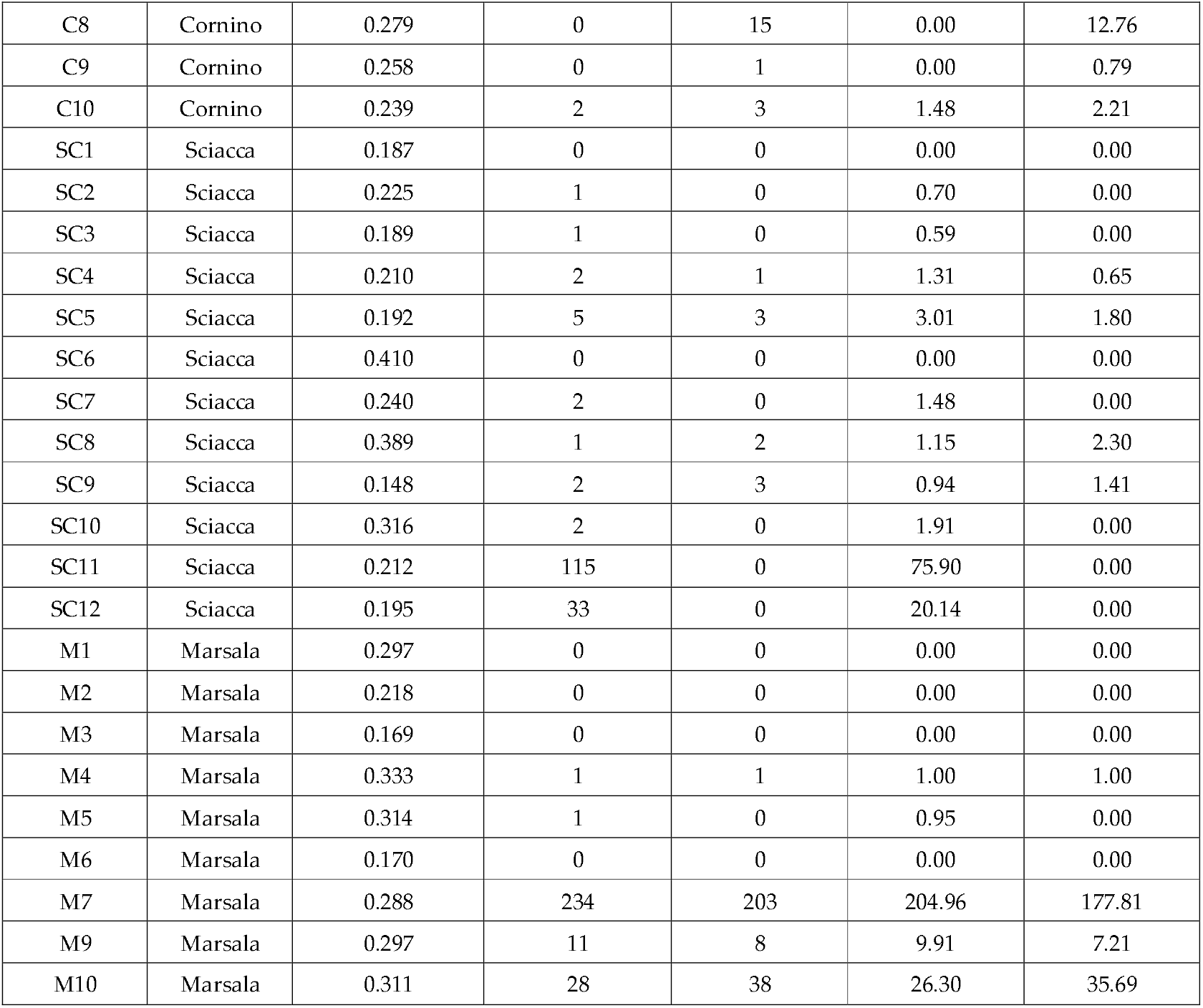

| Seed name | Location | Seed weight (g) | Total microbial colonies counted<br>(plate with dilution zero) |  | CFU/g |  |
| --- | --- | --- | --- | --- | --- | --- |
|  |  |  | SGY | NA | SGY | NA |
| SN1 | San Nicola | 0.327 | 301 | 202 | 295.84 | 198.54 |
| SN2 | San Nicola | 0.205 | 104 | 6 | 66.52 | 3.84 |
| SN3 | San Nicola | 0.174 | 0 | 0 | 0.00 | 0.00 |
| SN4 | San Nicola | 0.312 | 0 | 0 | 0.00 | 0.00 |
| SN5 | San Nicola | 0.293 | 0 | 0 | 0.00 | 0.00 |
| SN6 | San Nicola | 0.054 | 2 | 14 | 0.35 | 2.48 |
| SN7 | San Nicola | 0.211 | 102 | 87 | 67.03 | 57.17 |
| SN8 | San Nicola | 0.363 | 0 | 0 | 0.00 | 0.00 |
| E1 | Erice | 0.220 | 1 | 0 | 0.68 | 0.00 |
| E2 | Erice | 0.266 | 2 | 0 | 1.63 | 0.00 |
| E3 | Erice | 0.182 | 0 | 0 | 0.00 | 0.00 |
| E4 | Erice | 0.286 | 35 | 30 | 30.46 | 26.11 |
| E5 | Erice | 0.089 | 7 | 5 | 2.02 | 1.44 |
| C1 | Cornino | 0.320 | 376 | 352 | 362.41 | 339.28 |
| C2 | Cornino | 0.292 | 47 | 3 | 41.69 | 2.66 |
| C3 | Cornino | 0.145 | 248 | 260 | 114.34 | 119.87 |
| C4 | Cornino | 0.218 | 6 | 0 | 4.06 | 0.00 |
| C5 | Cornino | 0.186 | 7 | 0 | 4.09 | 0.00 |
| C6 | Cornino | 0.163 | 300 | 300 | 154.60 | 154.60 |
| C7 | Cornino | 0.217 | 0 | 0 | 0.00 | 0.00 |
| C8 | Cornino | 0.279 | 0 | 15 | 0.00 | 12.76 |
| C9 | Cornino | 0.258 | 0 | 1 | 0.00 | 0.79 |
| C10 | Cornino | 0.239 | 2 | 3 | 1.48 | 2.21 |
| SC1 | Sciacca | 0.187 | 0 | 0 | 0.00 | 0.00 |
| SC2 | Sciacca | 0.225 | 1 | 0 | 0.70 | 0.00 |
| SC3 | Sciacca | 0.189 | 1 | 0 | 0.59 | 0.00 |
| SC4 | Sciacca | 0.210 | 2 | 1 | 1.31 | 0.65 |
| SC5 | Sciacca | 0.192 | 5 | 3 | 3.01 | 1.80 |
| SC6 | Sciacca | 0.410 | 0 | 0 | 0.00 | 0.00 |
| SC7 | Sciacca | 0.240 | 2 | 0 | 1.48 | 0.00 |
| SC8 | Sciacca | 0.389 | 1 | 2 | 1.15 | 2.30 |
| SC9 | Sciacca | 0.148 | 2 | 3 | 0.94 | 1.41 |
| SC10 | Sciacca | 0.316 | 2 | 0 | 1.91 | 0.00 |
| SC11 | Sciacca | 0.212 | 115 | 0 | 75.90 | 0.00 |
| SC12 | Sciacca | 0.195 | 33 | 0 | 20.14 | 0.00 |
| M1 | Marsala | 0.297 | 0 | 0 | 0.00 | 0.00 |
| M2 | Marsala | 0.218 | 0 | 0 | 0.00 | 0.00 |
| M3 | Marsala | 0.169 | 0 | 0 | 0.00 | 0.00 |
| M4 | Marsala | 0.333 | 1 | 1 | 1.00 | 1.00 |
| M5 | Marsala | 0.314 | 1 | 0 | 0.95 | 0.00 |
| M6 | Marsala | 0.170 | 0 | 0 | 0.00 | 0.00 |
| M7 | Marsala | 0.288 | 234 | 203 | 204.96 | 177.81 |
| M9 | Marsala | 0.297 | 11 | 8 | 9.91 | 7.21 |
| M10 | Marsala | 0.311 | 28 | 38 | 26.30 | 35.69 |

### Appendix 2 Morphological characterization of the 44 selected endophytes isolated from *Posidonia oceanica* seeds

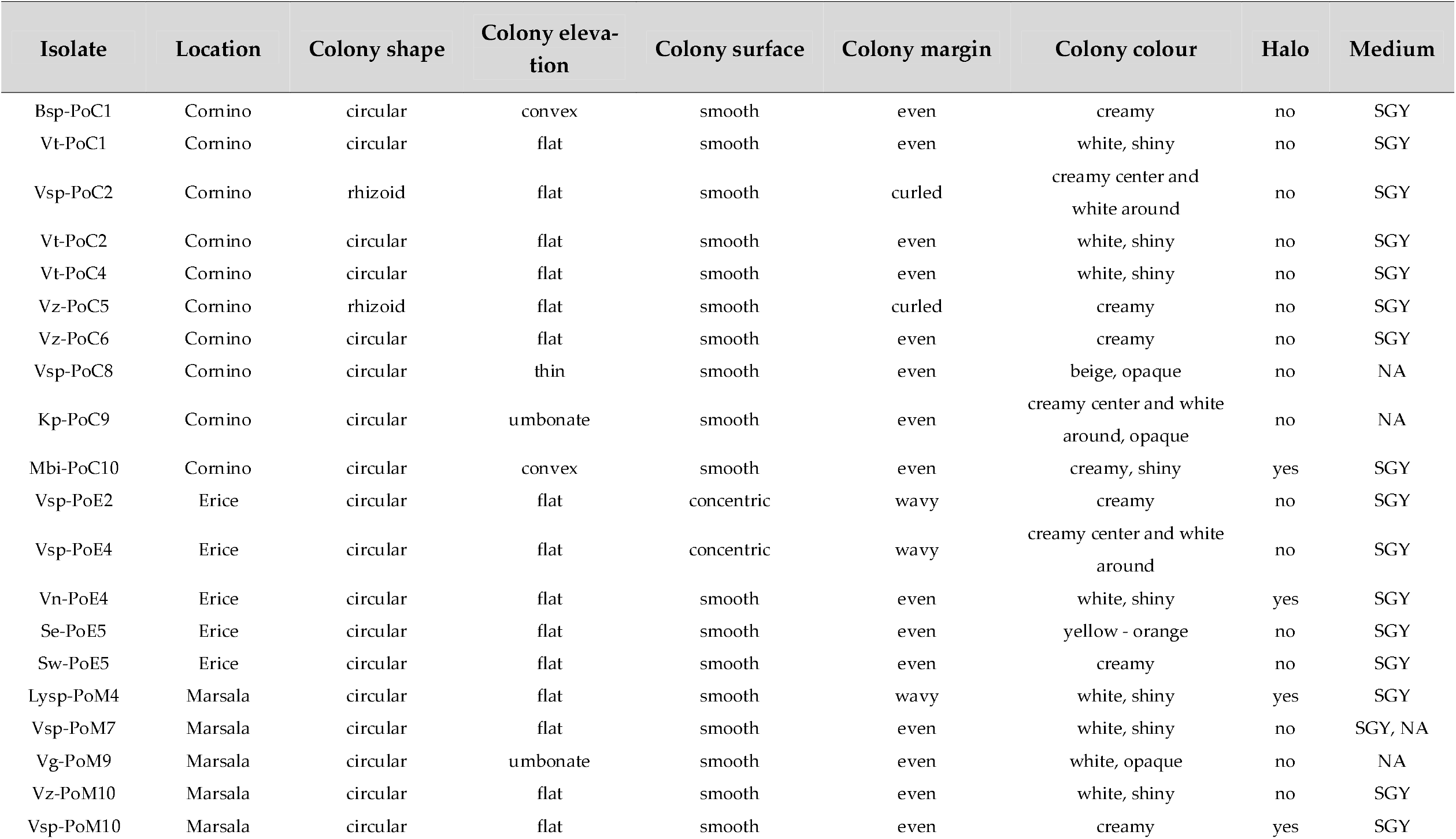
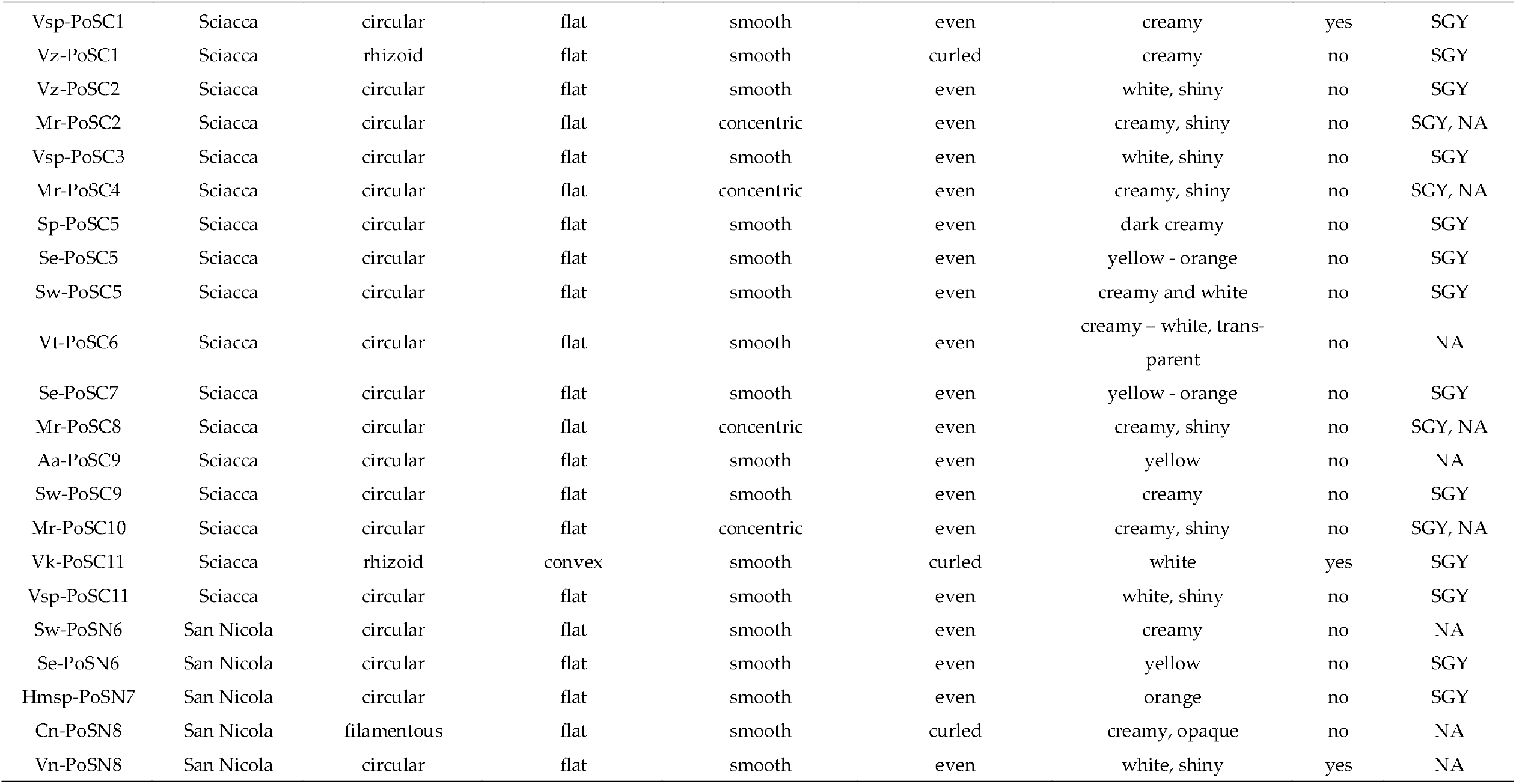
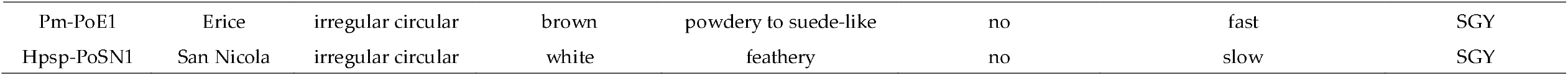

**Figure A1:**
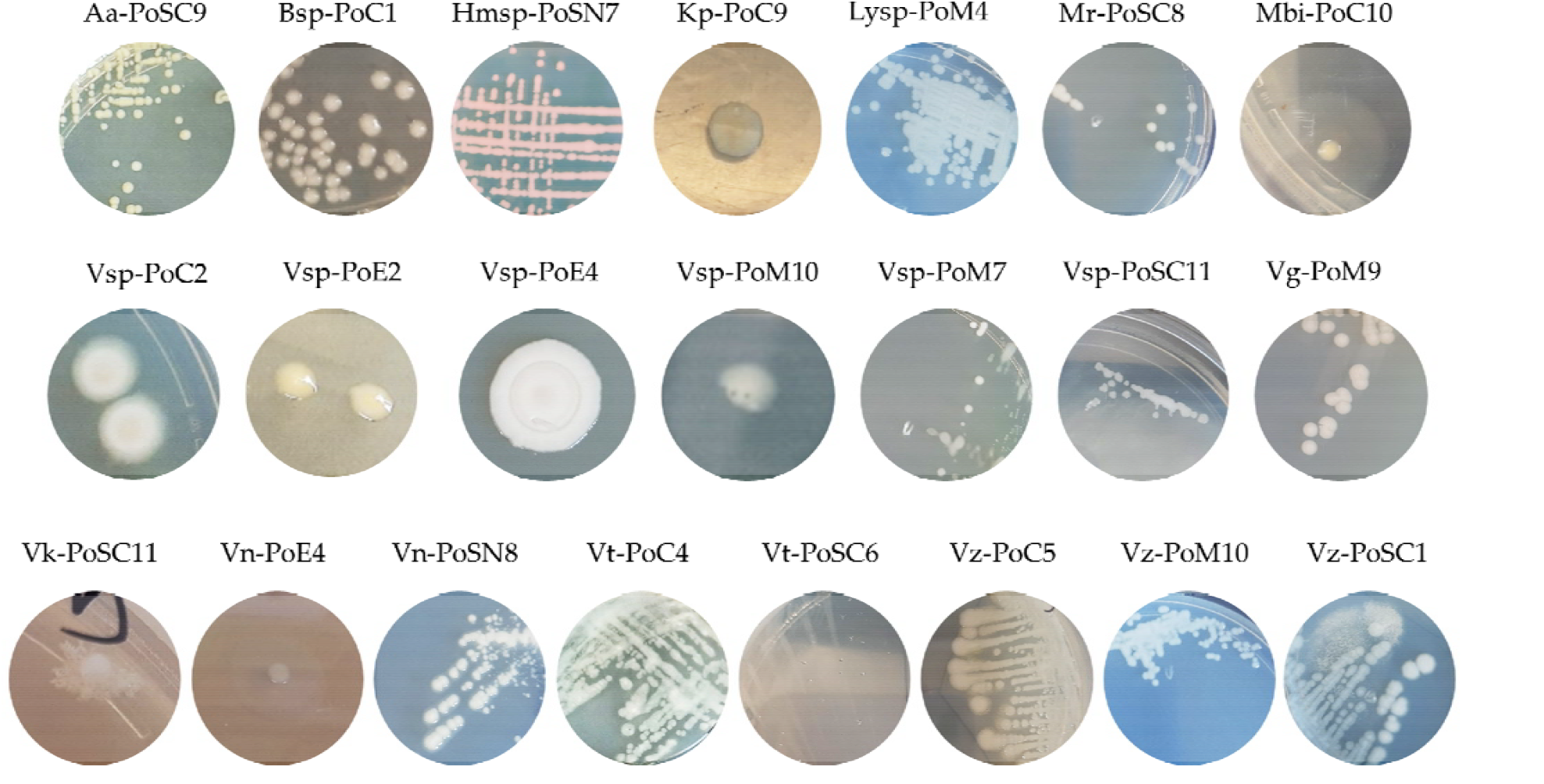
Morphology of 22 bacterial isolates characterized for plant growth-promoting properties.

## References

Alagna A, Giacalone VM, Zenone A, et al (2024) Tannins and copper sulphate as antimicrobial agents to prevent contamination of Posidonia oceanica seedling culture for restoration purposes. Front Plant Sci 15:1433358. 10.3389/fpls.2024.1433358

Ali B, Hafeez A, Javed MA, et al (2022) Role of endophytic bacteria in salinity stress amelioration by physiological and molecular mechanisms of defense: A comprehensive review. South African Journal of Botany 151:33–46. 10.1016/j.sajb.2022.09.036

Arnaud-Haond S, Migliaccio M, Diaz-Almela E, et al (2007) Vicariance patterns in the Mediterranean Sea: east–west cleavage and low dispersal in the endemic seagrass Posidonia oceanica. J Biogeogr 34:963–976. 10.1111/j.1365-2699.2006.01671.x

Bacci T, Scardi M, Tomasello A, Valiante LM, Piazzi L, Calvo S, Badalamenti F, Di nuzzo F, Raimondi V, Assenzo M, Cecchi E, Penna M, Bertasi F, Piazzi A, La Porta B (2025) Long-term response of Posidonia oceanica meadow restoration at the population and plant level: implications for management decisions. Restoration Ecology 33 (3):e14360. 1-12. 10.1111/rec.14360

Badalamenti F, Alagna A, D’Anna G, Terlizzi A, Di Carlo G (2011) The impact of dredge-fill on Posidonia oceanica seagrass Meadows: Regression and patterns of recovery. Marine Pollution Bulletin 62:483–489. 10.1016/j.marpolbul.2010.12.011

Bashan Y, Moreno M, Troyo E (2000) Growth promotion of the seawater-irrigated oilseed halophyte Salicornia bigelovii inoculated with mangrove rhizosphere bacteria and halotolerant Azospirillum spp. Biol Fertil Soils 32:265–272. 10.1007/s003740000246

Becker R, Ulrich K, Behrendt U, et al (2022) Genomic Characterization of Aureimonas altamirensis C2P003—A Specific Member of the Microbiome of Fraxinus excelsior Trees Tolerant to Ash Dieback. Plants 11:1–17. 10.3390/plants11243487

Bilal S, Khan AL, Shahzad R, et al (2017) Endophytic Paecilomyces formosus LHL10 Augments Glycine max L. Adaptation to Ni-contamination through affecting endogenous phytohormones and oxidative stress. Front Plant Sci 8:870, 1–17. 10.3389/fpls.2017.00870

Blanchet E, Prado S, Stien D, et al (2017) Quorum sensing and quorum quenching in the Mediterranean Seagrass Posidonia oceanica microbiota. Front Mar Sci 4:1–10. 10.3389/fmars.2017.00218

Brescia F, Pertot I, Puopolo G (2020) Lysobacter. In: Beneficial Microbes in Agro-Ecology: Bacteria and Fungi. Academic Press, 313–338

Calvo S, Calvo R, Luzzu F, et al (2021) Performance assessment of Posidonia oceanica (L.) Delile restoration experiment on dead matte twelve years after planting—structural and functional meadow features. Water 13:724, 1–17. 10.3390/w13050724

Calvo S, Tomasello A, Di Maida G, et al (2010) Seagrasses along the Sicilian coasts. Chemistry and Ecology 26:249–266. 10.1080/02757541003636374

Castejón-Silvo I, Terrados J (2021) Poor success of seagrass Posidonia oceanica transplanting in a meadow disturbed by power line burial. Mar Environ Res 170:1–10. 10.1016/j.marenvres.2021.105406

Celdrán D, Espinosa E, Sánchez-Amat A, Marín A (2012) Effects of epibiotic bacteria on leaf growth and epiphytes of the seagrass Posidonia oceanica. Mar Ecol Prog Ser 456:21–27. 10.3354/meps09672

Chaudhry V, Patil PB (2016) Genomic investigation reveals evolution and lifestyle adaptation of endophytic Staphylococcus epidermidis. Sci Rep 6:1–11. 10.1038/srep19263

Chaudhry V, Sharma S, Bansal K, Patil PB (2017) Glimpse into the genomes of rice endophytic bacteria: Diversity and distribution of Firmicutes. Front Microbiol 7:2115. 10.3389/fmicb.2016.02115

Chiellini C, Maida I, Emiliani G, et al (2015) Endophytic and rhizospheric bacterial communities isolated from the medicinal plants Echinacea purpurea and Echinacea angustifolia. International Microbiology 17:165–174. 10.2436/20.1501.01.219

Compant S, Brader G, Muzammil S, et al (2013) Use of beneficial bacteria and their secondary metabolites to control grapevine pathogen diseases. BioControl 58:435–455. 10.1007/s10526-012-9479-6

Compant S, Samad A, Faist H, Sessitsch A (2019) A review on the plant microbiome: Ecology, functions, and emerging trends in microbial application. J Adv Res 19:29–37. 10.1016/j.jare.2019.03.004

Cramer MJ, Haghshenas N, Bagwell CE, et al (2011) Celerinatantimonas diazotrophica gen. nov., sp. nov., a nitrogen-fixing bacterium representing a new family in the Gammaproteobacteria, Celerinatantimonadaceae fam. nov. Int J Syst Evol Microbiol 61:1053–1060. 10.1099/ijs.0.017905-0

da Silva N, Taniwaki MH, Junqueria VCA, et al (2013) Microbiological examination methods of food and water: a laboratory manual. 2nd Edition. CRC Press, 1–469. 10.1201/9781315165011

del Castillo I, Hernández P, Lafuente A, et al (2012) Self-bioremediation of cork-processing wastewaters by (chloro)phenol-degrading bacteria immobilised onto residual cork particles. Water Res 46:1723–1734. 10.1016/j.watres.2011.12.038

Di Carlo G, Badalamenti F, Jensen AC, et al (2005) Colonisation process of vegetative fragments of Posidonia oceanica (L.) Delile on rubble mounds. Mar Biol 147:1261–1270. 10.1007/s00227-005-0035-0

Döebereiner J (1995) Isolation and identification of aerobic nitrogen-fixing bacteria from soil and plants. In: Alef K, Nannipieri P (eds) Methods in Applied Soil Microbiology and Biochemistry. Academic Press, London, UK, 134–141

Doyle JJ, Doyle JL. (1987) A rapid DNA isolation procedure for small quantities of fresh leaf tissue. Phytochemical Bulletin 19:11–15

Drenker C, El Mazouar D, Bücker G, et al (2023) Characterization of a Disease-Suppressive Isolate of Lysobacter enzymogenes with Broad Antagonistic Activity against Bacterial, Oomycetal and Fungal Pathogens in Different Crops. 10.3390/plants12030682

Elbeltagy A, Nishioka K, Suzuki H, et al (2000) Isolation and characterization of endophytic bacteria from wild and traditionally cultivated rice varieties. Soil Sci Plant Nutr 46:617–629. 10.1080/00380768.2000.10409127

Espinosa E, Marco-Noales E, Gómez D, et al (2010) Taxonomic study of Marinomonas strains isolated from the seagrass Posidonia oceanica, with descriptions of Marinomonas balearica sp. nov. and Marinomonas pollencensis sp. nov. Int J Syst Evol Microbiol 60:93–98. 10.1099/ijs.0.008607-0

Faddetta T, Abbate L, Alibrandi P, et al (2021) The endophytic microbiota of Citrus limon is transmitted from seed to shoot highlighting differences of bacterial and fungal community structures. Sci Rep 11:1–12. 10.1038/s41598-021-86399-5

FAO (2024) Global status of salt-affected soils - Main report. FAO, Rome

Fidalgo C, Henriques I, Rocha J, et al (2016) Culturable endophytic bacteria from the salt marsh plant Halimione portulacoides: phylogenetic diversity, functional characterization, and influence of metal(loid) contamination. Environmental Science and Pollution Research 23:10200–10214. 10.1007/s11356-016-6208-1

Frank AC, Guzmán JPS, Shay JE (2017) Transmission of bacterial endophytes. Microorganisms 5:70, 1–21. 10.3390/microorganisms5040070

Gabriele M, Vitali F, Chelucci E, Chiellini C (2022) Characterization of the Cultivable Endophytic Bacterial Community of Seeds and Sprouts of Cannabis sativa L. and Perspectives for the Application as Biostimulants. Microorganisms 10:1742, 1–20. 10.3390/microorganisms10091742

Garcias-Bonet N, Arrieta JM, de Santana CN, et al (2012) Endophytic bacterial community of a Mediterranean marine angiosperm (Posidonia oceanica). Front Microbiol 3:342, 1–16. 10.3389/fmicb.2012.00342

Garcias-Bonet N, Arrieta JM, Duarte CM, Marbà N (2016) Nitrogen-fixing bacteria in Mediterranean seagrass (Posidonia oceanica) roots. Aquat Bot 131:57–60. 10.1016/j.aquabot.2016.03.002

Gordon SA, Weber RP (1951) Colorimetric Estimation of Indoleacetic Acid. Plant Physiol 26:192–195. 10.1104/pp.26.1.192

Govers LL, Man In ‘t Veld WA, Meffert JP, et al (2016) Marine Phytophthora species can hamper conservation and restoration of vegetated coastal ecosystems. Proceedings Royal Society B 283:20160812: 10.1098/rspb.2016.0812

Hardoim PR, van Overbeek LS, Berg G, et al (2015) The Hidden World within Plants: Ecological and Evolutionary Considerations for Defining Functioning of Microbial Endophytes. Microbiology and Molecular Biology Reviews 79:293–320. 10.1128/mmbr.00050-14

Harley JP, Prescott LM (2002) Laboratory Exercises in Microbiology, Fifth Edition. The McGraw-Hill Companies, New York, NY, USA

Higgins JJ (2004) Introduction to Modern Nonparametric Statistics. Brooks/Cole -Thomson Learning, United States of America

Houbraken J, Kocsubé S, Visagie CM, et al (2020) Classification of Aspergillus, Penicillium, Talaromyces and related genera (Eurotiales): An overview of families, genera, subgenera, sections, series and species. Stud Mycol 95:5–169. 10.1016/j.simyco.2020.05.002

Huang WS, Wang LT, Chen JS, et al (2021) Vibrio nitrifigilis sp. nov., a marine nitrogen-fixing bacterium isolated from the lagoon sediment of an islet inside an atoll. Antonie van Leeuwenhoek, International Journal of General and Molecular Microbiology 114:933–945. 10.1007/s10482-021-01567-x

Jiang C, Tanaka M, Nishikawa S, et al (2022) Vibrio Clade 3.0: New Vibrionaceae Evolutionary Units Using Genome-Based Approach. Curr Microbiol 79:1–15. 10.1007/s00284-021-02725-0

John JE, Maheswari M, Kalaiselvi T, et al (2023) Biomining Sesuvium portulacastrum for halotolerant PGPR and endophytes for promotion of salt tolerance in Vigna mungo L. Front Microbiol 14:1–18. 10.3389/fmicb.2023.1085787

Kearl J, McNary C, Lowman JS, et al (2019) Salt-tolerant halophyte rhizosphere bacteria stimulate growth of alfalfa in salty soil. Front Microbiol 10:1–11. 10.3389/fmicb.2019.01849

Khare E, Mishra J, Arora NK (2018) Multifaceted interactions between endophytes and plant: Developments and Prospects. Front Microbiol 9:1–12. 10.3389/fmicb.2018.02732

Kiki MJ (2016) A new medium for the isolation and enrichment of halophilic actinobacteria. Life Sci J 13:65–71. 10.7537/marslsj13011610

Lane DJ (1991) 16S/23S rRNA sequencing. Nucleic acid techniques in bacterial systematics.

Lee H, Golicz AA, Bayer PE, et al (2018) Genomic comparison of two independent seagrass lineages reveals habitat-driven convergent evolution. J Exp Bot 69:3689–3702. 10.1093/jxb/ery147

Liu BB, Wang HF, Li QL, et al (2016) Aurantimonas endophytica sp. nov., A novel endophytic bacterium isolated from roots of Anabasis elatior (C. A. Mey.) Schischk. Int J Syst Evol Microbiol 66:4112–4117. 10.1099/ijsem.0.001320

Liu Y, Zuo S, Zou Y, et al (2013) Investigation on diversity and population succession dynamics of endophytic bacteria from seeds of maize (Zea mays L., Nongda108) at different growth stages. Ann Microbiol 63:71–79. 10.1007/s13213-012-0446-3

Lopes R, Tsui S, Gonçalves PJRO, de Queiroz MV (2018) A look into a multifunctional toolbox: endophytic Bacillus species provide broad and underexploited benefits for plants. World J Microbiol Biotechnol 34:1–10. 10.1007/s11274-018-2479-7

Lucas-Elió P, Marco-Noales E, Espinosa E, et al (2011) Marinomonas alcarazii sp. nov., M. rhizomae sp. nov., M. foliarum sp. nov., M. posidonica sp. nov. and M. aquiplantarum sp. nov., isolated from the microbiota of the seagrass Posidonia oceanica. Int J Syst Evol Microbiol 61:2191–2196. 10.1099/ijs.0.027227-0

Maia C, Horta Jung M, Carella G, et al (2022) Eight new Halophytophthora species from marine and brackish-water ecosystems in Portugal and an updated phylogeny for the genus. Persoonia: Molecular Phylogeny and Evolution of Fungi 48:54–90. 10.3767/persoonia.2022.48.02

MarbàN, DCM (1998) Rhizome elongation and seagrass clonal growth. Mar Ecol Prog Ser 174:269–280

Mesa J, Mateos-Naranjo E, Caviedes MA, et al (2015) Scouting contaminated estuaries: Heavy metal resistant and plant growth promoting rhizobacteria in the native metal rhizoaccumulator Spartina maritima. Mar Pollut Bull 90:150–159. 10.1016/j.marpolbul.2014.11.002

Mohr W, Lehnen N, Ahmerkamp S, et al (2021) Terrestrial-type nitrogen-fixing symbiosis between seagrass and a marine bacterium. Nature 600:105–109. 10.1038/s41586-021-04063-4

Moreno-Gavíra A, Huertas V, Diánez F, et al (2020) Paecilomyces and its importance in the biological control of agricultural pests and diseases. Plants 9:1746, 1–28. 10.3390/plants9121746

Nautiyal CS (1999) An efficient microbiological growth medium for screening phosphate solubilizing microorganisms. FEMS Microbiol Lett 170:265–270. 10.1111/j.1574-6968.1999.tb13383.x

Nelson EB (2018) The seed microbiome: Origins, interactions, and impacts. Plant Soil 422:7–34. 10.1007/s11104-017-3289-7

Nguyen NH, Song Z, Bates ST, et al (2016) FUNGuild: An open annotation tool for parsing fungal community datasets by ecological guild. Fungal Ecol 20:241–248. 10.1016/J.FUNECO.2015.06.006

Ozan GN, Yılmaz F, Çaplık D, et al (2022) First report of pistachio die-back and canker disease caused by Paecilomyces maximus in Turkey. Journal of Plant Pathology 104:1165. 10.1007/s42161-022-01142-x

Pacifico D, Squartini A, Crucitti D, et al (2019) The Role of the Endophytic Microbiome in the Grapevine Response to Environmental Triggers. Front Plant Sci 10:1256, 1–15. 10.3389/fpls.2019.01256

Pathirana R, Carimi F (2024) Plant Biotechnology—An Indispensable Tool for Crop Improvement. Plants 13 (8): 1133, 1-18. 10.3390/plants13081133

Penrose DM, Glick BR (2003) Methods for isolating and characterizing ACC deaminase-containing plant growth-promoting rhizobacteria. Physiol Plant 118:10–15

Reusch TBH, Schubert PR, Marten SM, et al (2021) Lower Vibrio spp. abundances in Zostera marina leaf canopies suggest a novel ecosystem function for temperate seagrass beds. Mar Biol 168:1–6. 10.1007/s00227-021-03963-3

Román-Ponce B, Ramos-Garza J, Vásquez-Murrieta MS, et al (2016) Cultivable endophytic bacteria from heavy metal(loid)-tolerant plants. Arch Microbiol 198:941–956. 10.1007/s00203-016-1252-2

Rungjindamai N, Jones EBG (2024) Why Are There So Few Basidiomycota and Basal Fungi as Endophytes? A Review. Journal of Fungi 10:67, 1-30. 10.3390/jof10010067

Ruocco M, Lacorata G, Palatella L, Provera I, Zenone A, Martinez M, Datolo E, Pazzaglia J, Giacalone VM, Badalamenti F, Procaccini G (2024) Movement Ecology of a Coastal Foundation Seagrass Species: Insights From Genetic Data and Oceanographic Modelling. Diversity and Distributions 31:e13944. 1-18. 10.1111/ddi.13944

Sabernasab M, Jamali S, Marefat A, Abbasi S (2019) Molecular and Pathogenic Characteristics of Paecilomyces formosus, a New Causal Agent of Oak Tree Dieback in Iran. Forest Science 65:743–750. 10.1093/forsci/fxz045

Sampaio A, Silva V, Poeta P, Aonofriesei F (2022) Vibrio spp.: Life Strategies, Ecology, and Risks in a Changing Environment. Diversity (Basel) 14:1–26. 10.3390/d14020097

Santoyo G, Moreno-Hagelsieb G, del Carmen Orozco-Mosqueda M, Glick BR (2016) Plant growth-promoting bacterial endophytes. Microbiol Res 183:92–99. 10.1016/j.micres.2015.11.008

Scanu S, Piazzolla D, Bonamano S, et al (2022) Economic Evaluation of Posidonia oceanica Ecosystem Services along the Italian Coast. Sustainability (Switzerland) 14:1–17. 10.3390/su14010489

Schwyn B, Neilands JB (1987) Universal chemical assay for the detection and determination of siderophores. Anal Biochem 160:47–56. 10.1016/0003-2697(87)90612-9

Setiawan A, Setiawan F, Juliasih NLGR, et al (2022) Fungicide Activity of Culture Extract from Kocuria palustris 19C38A1 against Fusarium oxysporum. Journal of Fungi 8:1–11. 10.3390/jof8030280

Shurigin V, Egamberdieva D, Li L, et al (2020) Endophytic bacteria associated with halophyte Seidlitzia rosmarinus Ehrenb. ex Boiss. from saline soil of Uzbekistan and their plant beneficial traits. J Arid Land 12:730–740. 10.1007/s40333-020-0019-4

Stackebrandt E, Mondotte JA, Fazio LL, Jetten M (2022) Authors need to be prudent when assigning names to microbial isolates. Antonie van Leeuwenhoek, International Journal of General and Molecular Microbiology 115:1–5. 10.1007/s10482-021-01675-8

Surette MA, Sturz A V, Lada RR, Nowak J (2003) Bacterial endophytes in processing carrots (Daucus carota L. var. sativus): their localization, population density, biodiversity and their effects on plant growth. Plant and Soil 253:381–390

Sutera A, Bonaviri C, Spinelli P, et al (2024) Fruit encasing preserves the dispersal potential and viability of stranded Posidonia oceanica seeds. Sci Rep 14:6218. 10.1038/s41598-024-56536-x

Sutera A, Spinelli P, Pacifico D, et al (2025) Extended storage of Posidonia oceanica seeds and seedlings: A breakthrough for year-round seagrass restoration. Biol Conserv 310:111385. 10.1016/j.biocon.2025.111385

Tarquinio F, Attlan O, Vanderklift MA, et al (2021) Distinct Endophytic Bacterial Communities Inhabiting Seagrass Seeds. Front Microbiol 12:1–17. 10.3389/fmicb.2021.703014

Tarquinio F, Hyndes GA, Laverock B, et al (2019) The seagrass holobiont: Understanding seagrass-bacteria interactions and their role in seagrass ecosystem functioning. FEMS Microbiol Lett 366:fnz057. 10.1093/femsle/fnz057

Telesca L, Belluscio A, Criscoli A, et al (2015) Seagrass meadows (Posidonia oceanica) distribution and trajectories of change. Sci Rep 5:1–14. 10.1038/srep12505

Torta L, Burruano S, Giambra S, et al (2022) Cultivable Fungal Endophytes in Roots, Rhizomes and Leaves of Posidonia oceanica (L.) Delile along the Coast of Sicily, Italy. Plants 11:1139. 10.3390/plants11091139

Torta L, Lo Piccolo S, Piazza G, et al (2015) Lulwoana sp., a dark septate endophyte in roots of Posidonia oceanica (L.) Delile seagrass. Plant Biology 17:505–511. 10.1111/plb.12246

Truyens S, Weyens N, Cuypers A, Vangronsveld J (2015) Bacterial seed endophytes: Genera, vertical transmission and interaction with plants. Environ Microbiol Rep, 7:1–11. 10.1111/1758-2229.12181

Tu C-K, Wang P-H, Lee M-H (2023) The endophytic bacterium Lysobacter firmicutimachus strain 5-7 is a promising biocontrol agent against rice seedling disease caused by Pythium arrhenomanes in nursery trays. Plant Disease 107:1075–1086. 10.1094/PDIS-05-22-1195-RE

Ugarelli K, Chakrabarti S, Laas P, Stingl U (2017) The seagrass holobiont and its microbiome. Microorganisms 5:81, 1–28. 10.3390/microorganisms5040081

Vilgalys R, Hester M (1990) Rapid Genetic Identification and Mapping of Enzymatically Amplified Ribosomal DNA from Several Cryptococcus Species. Journal of Bacteriology 172:8, 4238-4246. 10.1128/jb.172.8.4238-4246.1990

Vohník M (2022) Are lulworthioid fungi dark septate endophytes of the dominant Mediterranean seagrass Posidonia oceanica? Plant Biology 24:127–133. 10.1111/plb.13353

Vohník M, Borovec O, Kolařík M (2016) Communities of Cultivable Root Mycobionts of the Seagrass Posidonia oceanica in the Northwest Mediterranean Sea Are Dominated by a Hitherto Undescribed Pleosporalean Dark Septate Endophyte. Microb Ecol 71:442–451. 10.1007/s00248-015-0640-5

Vohník M, Borovec O, Kolaříková Z, et al (2019) Extensive sampling and high-throughput sequencing reveal Posidoniomyces atricolor gen. Et sp. Nov. (Aigialaceae, Pleosporales) as the dominant root mycobiont of the dominant Mediterranean seagrass Posidonia oceanica. MycoKeys 55:59–86. 10.3897/mycokeys.55.35682

Wang H, Narsing Rao MP, Gao Y, et al (2021) Insights into the endophytic bacterial community comparison and their potential role in the dimorphic seeds of halophyte Suaeda glauca. BMC Microbiol 21:1–16. 10.1186/s12866-021-02206-1

White TJ, Bruns T, Lee S, et al (1990) Amplification and direct sequencing of fungal ribosomal RNA Genes for phylogenetics. In: PCR Protocols: A Guide to Methods and Applications. Accademic Press Inc., 315–322

Wilson K (1997) Preparation of Genomic DNA from Bacteria. Current Protocols in Molecular Biology 56:2.4.1-2.4.5

Yin Z, Wang X, Hu Y, et al (2022) Metabacillus dongyingensis sp. nov. Is Represented by the Plant Growth-Promoting Bacterium BY2G20 Isolated from Saline-Alkaline Soil and Enhances the Growth of Zea mays L. under Salt Stress. mSystems 7:1–16. 10.1128/msystems.01426-21

Zacaria Vital T, Román-Ponce B, Rivera Orduña FN, et al (2019) An endophytic Kocuria palustris strain harboring multiple arsenate reductase genes. Arch Microbiol 201:1285–1293. 10.1007/s00203-019-01692-2

Zhang J, Wang P, Tian H, et al (2018) Identification of interior salt-tolerant bacteria from ice plant Mesembryanthemum crystallinum and evaluation of their promoting effects. Symbiosis 76:243–252. 10.1007/s13199-018-0551-6

Zhang J, Wang P, Tian H, et al (2020) Transcriptome analysis of ice plant growth-promoting endophytic bacterium Halomonas sp. Strain MC1 to identify the genes involved in salt tolerance. Microorganisms 8:88, 1–18. 10.3390/microorganisms8010088

